# Distribution and restoration of serotonin-immunoreactive paraneuronal cells during caudal fin regeneration in zebrafish

**DOI:** 10.1101/698019

**Authors:** Désirée König, Paule Dagenais, Anita Senk, Valentin Djonov, Christof M. Aegerter, Anna Jaźwińska

**Author notes:** corresponding author: Address: Department of Biology, University of Fribourg, Chemin du Musée 10, 1700 Fribourg, Switzerland.

## Abstract

Aquatic vertebrates possess diverse types of sensory cells in their skin to detect stimuli in the water. In the adult zebrafish, a common model organism, the presence of such cells in fins has only rarely been studied. Here, we identified scattered serotonin (5-HT)-positive cells in the epidermis of the caudal fin. These cells were distinct from keratinocytes as revealed by their low immunoreactivity for cytokeratin and desmosome markers. Instead, they were detected by Calretinin (Calbindin-2) and Synaptic vesicle glycoprotein 2 (SV2) antibodies, indicating a calcium-regulated neurosecretory activity. Consistently, electron microscopy revealed abundant secretory organelles in desmosome-negative cells in the fin epidermis. Based on the markers, 5-HT, Calretinin and SV2, we referred to these cells as HCS-cells. We found that HCS-cells were spread throughout the entire caudal fin at an average density of 140 cells per mm^2^ on each fin surface. These cells were strongly enriched at ray bifurcations in wild type fins, as well as in elongated fins of *another longfin* mutant fish. To determine whether hydrodynamics play a role in the distribution of HCS-cells, we used an interdisciplinary approach and performed kinematic analysis. Measurements of particle velocity with a fin model revealed differences in fluid velocities between bifurcated rods and adjacent non-bifurcated regions. Therefore the accumulation of HCS-cells near bone bifurcations may be a biological adaptation for sensing of water parameters. The significance of this HCS-cell pattern is reinforced by the fact, that it is reestablished in the regenerated fin after amputation. Regeneration of HCS-cells was not impaired by the chemical inhibition of serotonin synthesis, suggesting that this neurotransmitter is not essential for the restorative process. In conclusion, our study identified a specific population of solitary paraneurons in the zebrafish fin, whose distribution correlates with fluid dynamics.

## Introduction

The skin is continuously exposed to environmental fluctuations and external signals. In the epidermis, specialized cells recognize specific stimuli and transmit this information to the body through secretion of active compounds, such as hormones and neurotransmitters (Slominski et al., 2012). The released messengers can act locally in a paracrine manner and remotely through humoral, immune and neural pathways. To respond appropriately, sensory structures evolved distributions suitable for the types of stimuli and the environmental media.

Aquatic vertebrates, such as lampreys, fish, and amphibian tadpoles, can detect hydrodynamic parameters and dissolved chemicals through their epidermal sensors adapted for underwater conditions (Atema et al., 1988; von der Emde et al., 2004; Collin et al., 2008; Finger, 2009). Their mucus-covered epidermis contains various types of specialized chemo- and mechanoreceptors. In teleost fish, epidermal sensors can be arranged into multicellular structures, *e.g.* taste buds and mechanoreceptive neuromasts, or they can occur as scattered cells, *e.g.* solitary chemosensory cells and Merkel-like cells (Whitear, 1992; Kotrschal, 1996; Kasumyan, 2011; Coombs et al., 2013).

According to a concept introduced by Fujita (Fujita et al., 1988), diverse types of skin sensory cells can be classified as paraneurons. This term describes neurosecretory cells located in epithelia and characterized by the presence of vesicles with neurotransmitters or other messengers, which can be released in response to adequate stimuli (Fujita, 1989). In fish, various types of solitary paraneurons have been reported to contain serotonin (5-hydroxytryptamine, 5-HT) (Zaccone et al., 1994). In larval zebrafish, this neurotransmitter has been detected in Merkel-like basal cells of the taste buds and cutaneous scattered neuroepithelial cells (Coccimiglio and Jonz, 2012; Zachar and Jonz, 2012; Soulika et al., 2016). In adult zebrafish, superficial serotonin-positive cells have been observed in the epidermis of the caudal fin, but they have not been further analyzed (Tornini et al., 2017).

The zebrafish caudal fin is a non-muscularized appendage. It is a commonly used model for studying vertebrate limb regeneration due to its easy accessibility and fast regeneration after amputation (Akimenko et al., 2003; Tornini and Poss, 2014; Pfefferli and Jaźwińska, 2015; Wehner and Weidinger, 2015). The caudal fin has a bi-lobed shape with a central cleft (Fig. 1A, B), which is supported by 16-18 primary bony rays. Most primary rays bifurcate one to three times along their length, allowing for the widening of the appendage; only the lateral-most rays and sometimes the medial ray do not bifurcate (Fig. 1B). The rays contain bilateral parenthetical bones, called hemirays or lepidotrichia, which are regularly segmented (Fig. 1C). The hemirays surround mesenchymal tissue composed of fibroblasts, collagenous matrix, blood vessels and nerves (Fig. 1D). The rays are spanned by softer tissue called interray, which also contains mesenchymal tissue (Fig. 1E). Both surfaces of the fin are covered in a multi-layered non-cornified epidermis.

**Figure 1.**
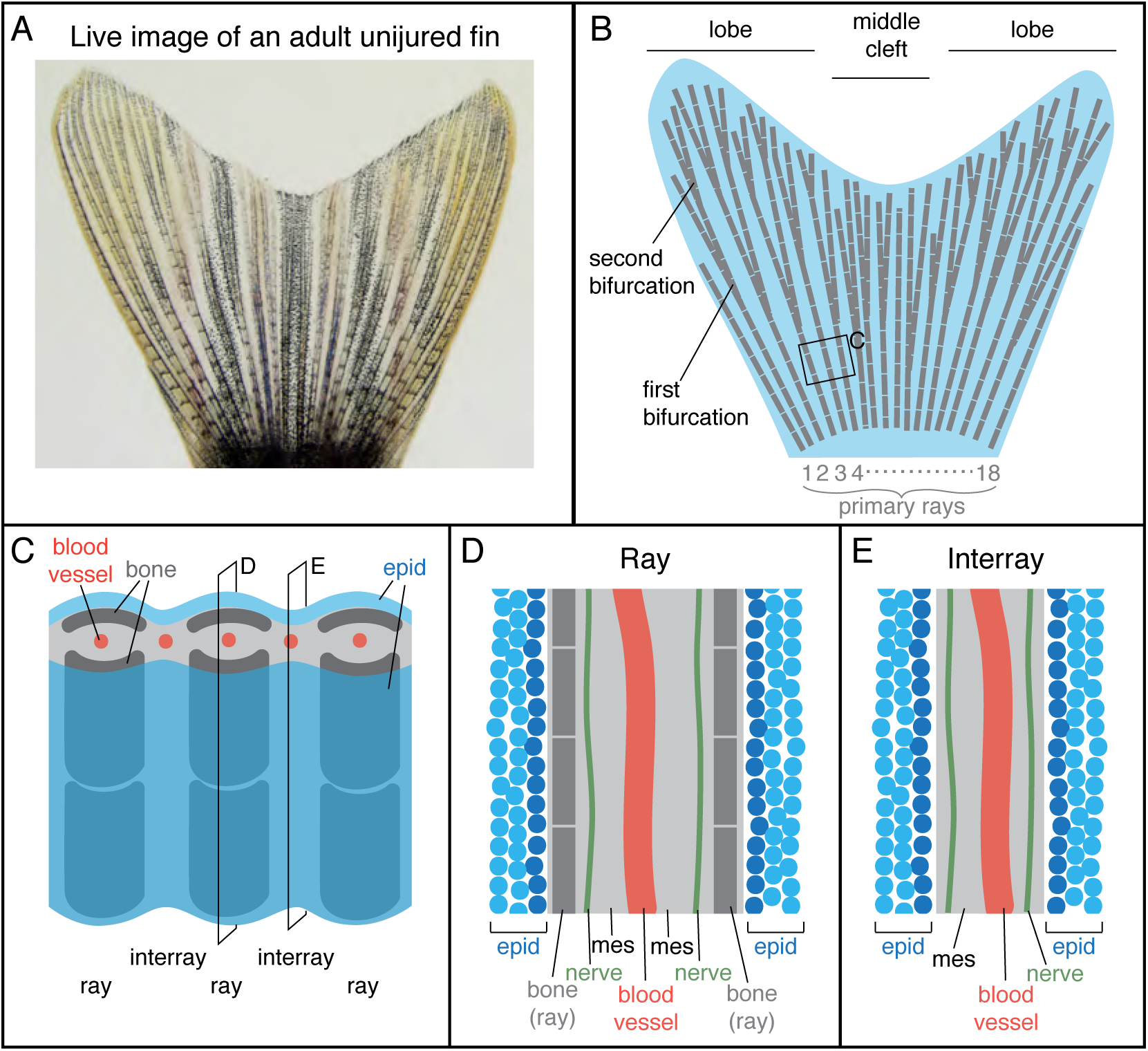
Schematic representation of an adult zebrafish caudal fin. (A) Live imaging of the caudal fin of an adult zebrafish. (B) Illustration shows the fan-like shape of the fin with longer lateral lobes and a shorter middle cleft. The fin fold contains 16-18 primary rays (gray) that are segmented and bifurcate once or twice. (C) Close-up representation of the fin shows a repetitive ray-interray arrangement. Rays are supported by bilateral dermal bones (dark gray). The fin is covered by epidermis (epid; blue) on both sides. The inner fin tissue contains mesenchyme (light gray) with centrally located blood vessels (red). (D-E) Longitudinal sections through a ray (D) and an interray (E) display a multilayered epidermis (blue) covering the fin surface. The mesenchyme (mes) is vascularized and innervated.

Experimental evidence can be found regarding the mechanosensory capacities of fins in various fish species (Flammang and Lauder, 2013; Williams Iv et al., 2013; Williams and Hale, 2015; Hardy et al., 2016; Aiello et al., 2017). The spatial distribution of mechanoreceptors across the skin is often correlated with regions that are optimally exposed to water flow, in order to maximize the chances of perceiving the cues (Collin et al., 2008; Bleckmann et al., 2014). Such strategic patterns depend on hydrodynamic properties of the fin surface that can be described by the principles of boundary layer, the layer at the surface of the body where the fluid is slowed or stationary. Specifically, the most important parameters are shear stress, acting parallel to the surface, and a velocity gradient, forming perpendicularly to the surface. Moreover, a turbulent boundary layer contains fluctuations of the fluid, such as eddies and swirls, which may be sensed by putative mechanical receptors inside the skin of an appendage. The presence of an excrescence, such as the bifurcating rays bulging out of the fin membrane, is likely to favor the breakout of turbulence. The advance of particle velocimetry imaging techniques provides new tools to quantify the boundary layer around the body (Anderson et al., 2001).

In this study, we aimed to characterized the morphology, the distribution and restoration of 5-HT-producing paraneurons in the adult caudal fin. We showed the expression of several molecular markers in these cells and their ultrastructure. Furthermore, we investigated these paraneurons during post-embryonic development in wild type fish and in adult mutants with elongated fins. Using a scaled ray model, we analyzed water flow velocities in the vicinity of bone bifurcations and correlated this with the distribution of 5-HT-positive cells. Finally, we performed fin amputation and characterize the regeneration of these cells. Overall, our study shows that small solitary 5-HT-immunoreactive cells represent a distinct population of paraneurons with putative mechanosensory functions.

## Results

### Identification of solitary 5-HT/Calretinin/SV2/-immunolabeled cells in the fin epidermis

The epidermis of the adult fin is mainly formed of 3-4 layers of keratinocytes (Fig. 1D, E). To investigate the presence of paraneuronal cells within this epidermis, we performed immunofluorescence analysis of longitudinal fin section using a rabbit antibody against 5-HT and a mouse antibody against Synaptic vesicle glycoprotein-2 (SV2). Both these markers labeled individual dispersed cells in the outer or sub-outer layer of the stratified epithelium (Fig. 2A). The shape of the 5-HT/SV2-positive cells was round at a diameter of 5.38 ± 0.61 μm (n = 7). High-resolution confocal imaging revealed that both molecules were distributed in a dotty pattern, suggesting a vesicular localization consistent with a neurosecretory function (Fig. 2A’). On confocal images, 5-HT and SV2 were often concentrated at one side of the cells, indicating their polarized nature. This polarization was not consistently oriented in one direction relative to the fin surface.

**Figure 2.**
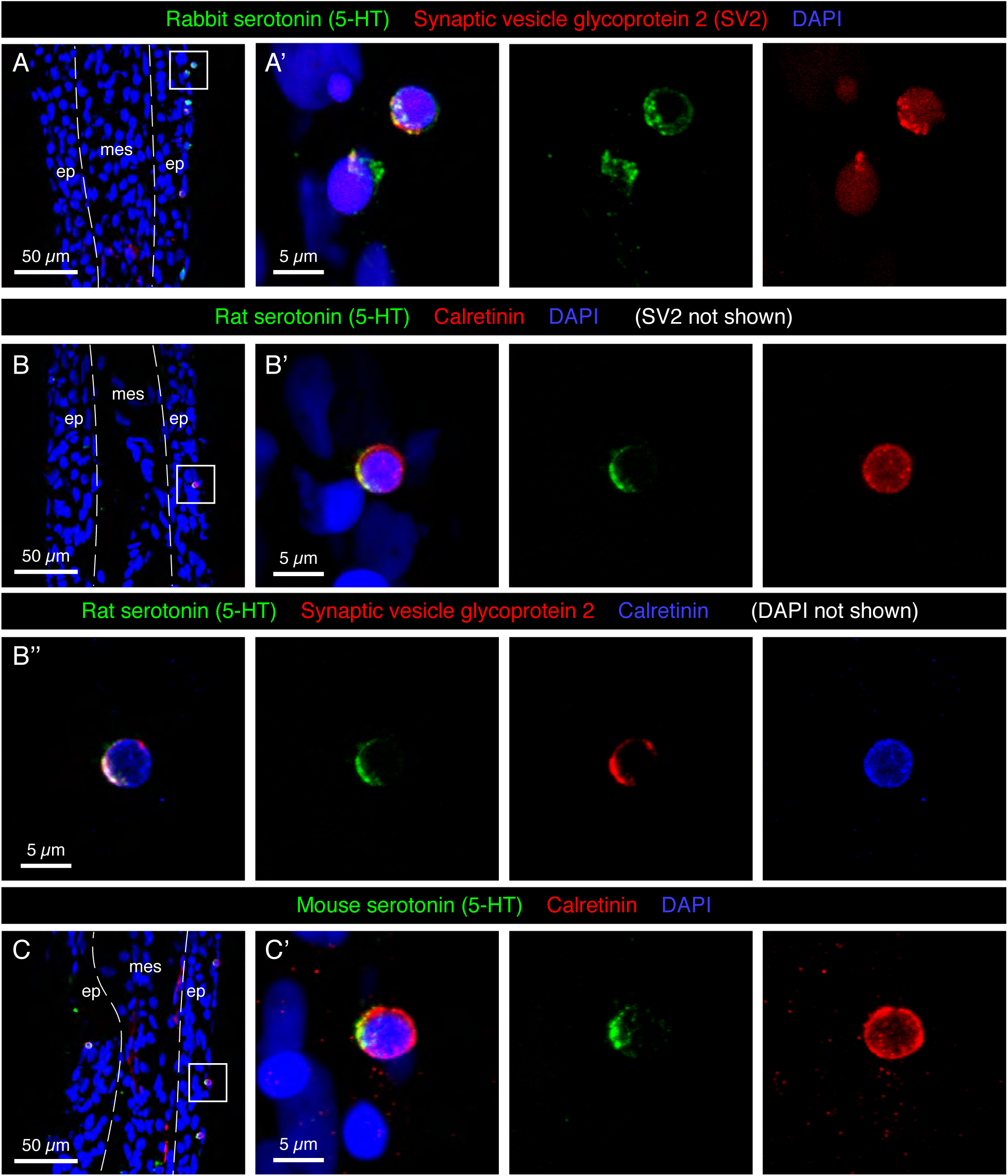
Molecular markers of HCS-cells in the adult fin epidermis. (A-C) Immunofluorescence staining of longitudinal sections of uninjured adult caudal fins; ep, epidermis, mes, mesenchyme. Dashed lines indicate the border between epidermis and mesenchyme. (A-A’) Rabbit antibody against serotonin (5-HT; green) and mouse antibody against Synaptic vesicle glycoprotein 2 (SV2; red) colocalize in single cells near the epidermal surface. (A’) A higher magnification of the framed area in (A) shows the vesicular and polarized distribution of both markers in the cells. (B-B’’) Triple immunostaining with rat antibody against serotonin (5-HT), rabbit antibody against Calretinin and mouse antibody against Synaptic vesicle glycoprotein 2. All three markers are expressed in the same cells of the epidermis. Serotonin and SV2 are polarized while Calretinin is found throughout the cytoplasm. (C-C’) Mouse antibody against serotonin and rabbit antibody against Calretinin label the same cells. All three serotonin antibodies have identical patterns. However, both Rat-serotonin and Mouse-serotonin required high concentrations and gave weaker signals than the rabbit antibody. N ≥ 4 for each staining. Nuclei are labeled with DAPI (blue). (A’, B’, C’) Images labeled with letters with prime symbols show higher magnifications of the frames in the corresponding images. The same rule applies to all the subsequent figures.

In certain fish species, such as sea catfish, appendages contain taste buds (Ikenaga and Kiyohara, 2018). In the developing zebrafish, taste buds contain one 5-HT-positive basal cell, which underlies Calretinin-expressing chemosensory cells (Zachar and Jonz, 2012; Soulika et al., 2016). To identify whether the 5-HT-positive cells of the epidermis are part of fin taste buds, we used a Calretinin antibody previously verified in zebrafish tissues by Western Blot and immunohistochemistry (Castro et al., 2006; Soulika et al., 2016). Accordingly, we performed a triple immunostaining with rat 5-HT, rabbit Calretinin and mouse SV2 antibodies. We found that all three markers were detected in the same solitary cells (Fig. 2B), and not in adjacent cells as in taste buds. To further validate this finding, we used another 5-HT antibody raised in mice. Consistently, we observed that both markers co-labeled the same small superficial cells (Fig. 2C). Unlike 5-HT, which appeared in vesicles, Calretinin displayed a non-polarized distribution in the cytoplasm. Calretinin expression was absent from other cells in the fin epidermis. The co-expression of a neurotransmitter, a Ca^2+^ buffering protein, and a synaptic vesicle glycoprotein suggests that the identified cells possess neurosecretory features. These molecular characteristics and the epidermal localization fit to the definition of paraneurons, according to the concept formulated by Fujita (Fujita, 1989). Based on the 5-HT, Calretinin, SV2 triple labeling, we abbreviated these markers and named this class of paraneurons HCS-cells.

### HCS-cells differ from keratinocytes

The intermediate filaments cytokeratins, are the most important structural elements of keratinocytes. To visualize the position of HCS-cells relative to keratinocytes, we performed immunofluorescence analysis using an antibody against cytokeratins. As previously shown (Jazwinska et al., 2007), keratin was detected in epidermal cells, which mostly had cubical or elongated spindle-like shapes (Fig. 3A). By contrast, the round 5-HT-labeled HCS-cells were devoid of keratins, suggesting a non-keratinocyte identity.

**Figure 3.**
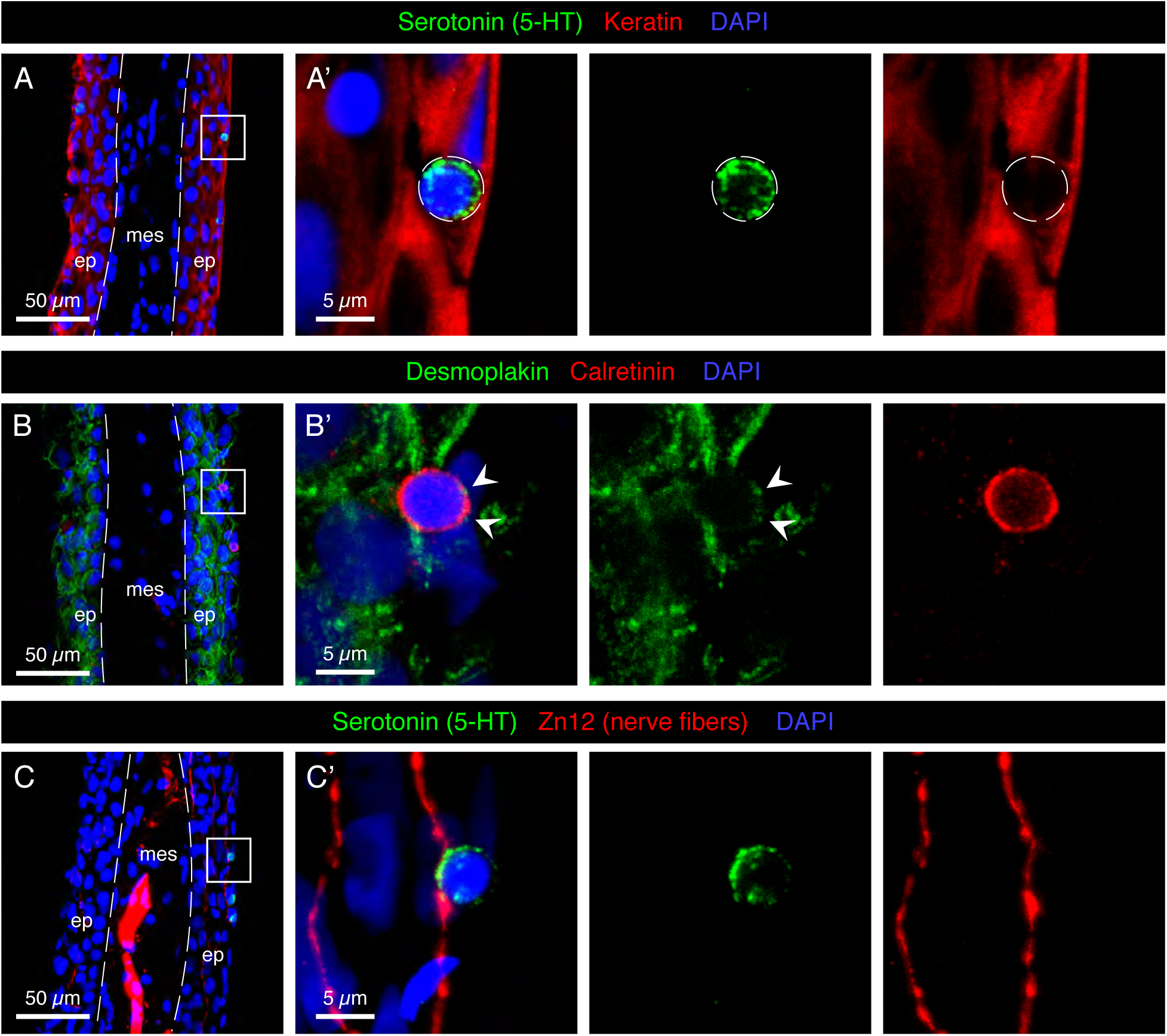
Characterization of HCS-cells in the adult uninjured fin. (A-C) Immunofluorescence staining of longitudinal sections of uninjured adult caudal fins; ep, epidermis, mes, mesenchyme. Dashed lines indicate the border between epidermis and mesenchyme. (A-A’) Serotonin-positive cells (green) do not express Keratin (red), which demarcates surrounding epidermal cells. (B-B’) Immunostaining for Desmoplakin (green) and Calretinin (red) indicates that Calretinin-positive cells possess very few desmosomes compared to the surrounding keratinocytes. Very weak dotty Desmoplakin staining is sometimes observed at the apical side of the cell (arrowheads). (C-C’) Staining for Serotonin (green) and Zn12 (red), a neuronal marker, reveals that cells are located close to nerve fibers. N ≥ 4 for each staining. Nuclei are labeled with DAPI (blue).

Keratin filaments are tethered to the plasma membrane by desmosomes, which form intercellular junctions. To determine whether HCS-cells form desmosomes, we visualized desmoplakin, a protein that links the cytoskeleton and the desmosomal plaque (Delva et al., 2009). Immunostaining with Desmoplakin 1/2 and 5-HT antibodies revealed that HCS-cells did not contain much of the desmosomal protein around their surface, in contrast to keratinocytes which were strongly outlined with this marker (Fig. 3B). Thus, HCS-cells are not connected to the surrounding epidermal cells through desmosomes.

Sensory cells typically form contacts with peripheral nerve fibers for signal transmission. To investigate whether the identified cells are innervated, we visualized neuronal processes with the Zn12 antibody (Metcalfe et al., 1990). Indeed, we found intraepidermal neural projections contacting 5-HT-positive cells (Fig. 3C). Thus, HCS-cells appear to be associated with the nervous system of the caudal fin.

### Ultrastructural features of paraneurons in the epidermis of the caudal fin

To determine the ultrastructure of HCS-cells, we performed transmission electron microscopy (EM) analysis on transversal sections of caudal fins. Tissue shrinkage is a common EM artifact caused by fixation, dehydration and embedding of tissues (Montesinos et al., 1983; Doughty et al., 1997). We found that processing for EM resulted in a global shrinkage of tissue nearly by a half compared to sections for confocal microscopy, but proportions between cells types were preserved. As previously reported (Iger and Wendelaar Bonga, 1994; Kotrschal et al., 1997), the surface of the epidermis was covered by squamous epithelial cells, referred to as the “pavement” cells (Fig. 4A, B). These cells are characterized by apical microplicae that form ridges on the cell surface, as shown by scanning EM (Fig. 4C). We searched for non-keratinocytes situated underneath this layer that would morphologically resemble the HCS-paraneurons observed by immunofluorescence.

**Figure 4.**
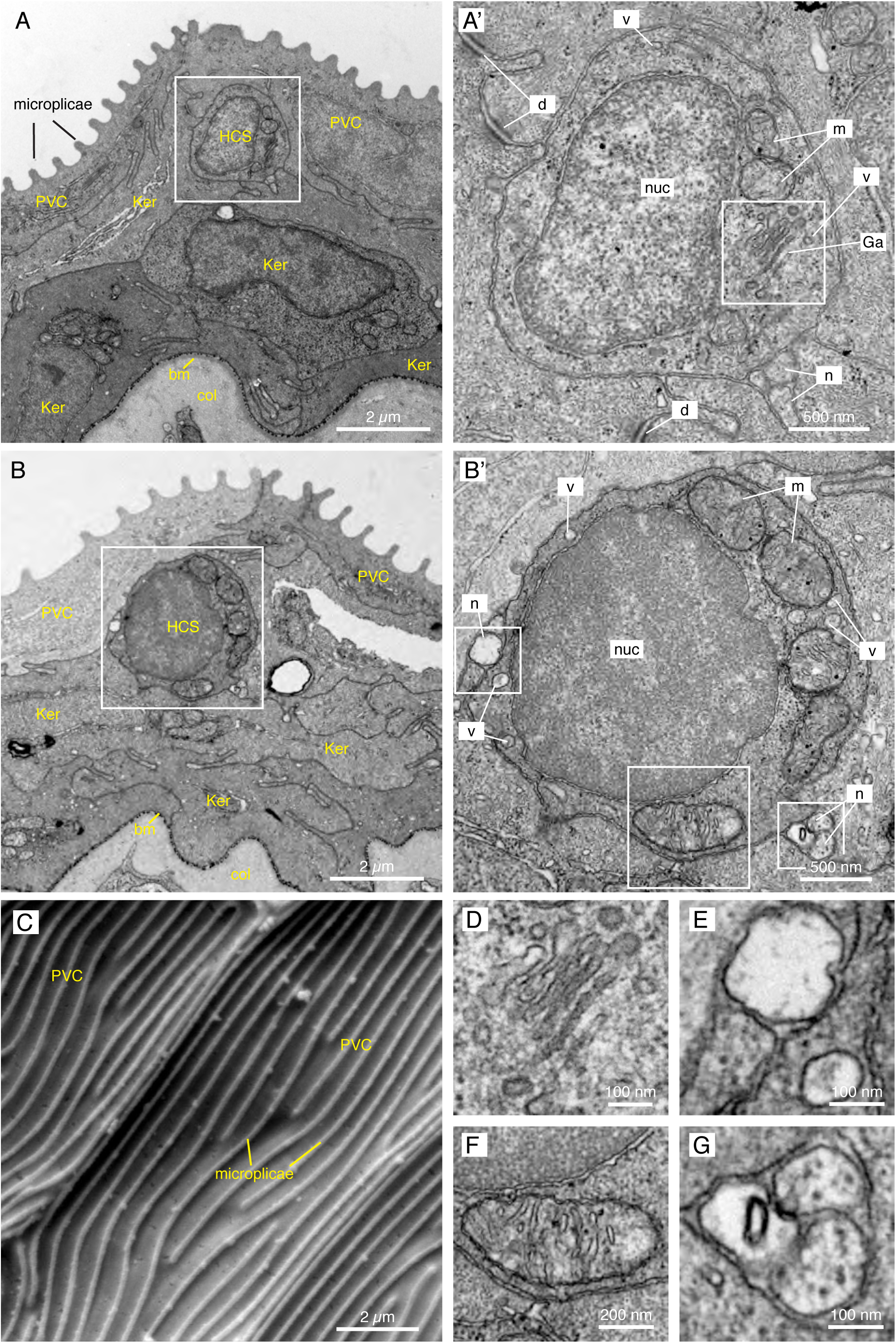
Electron microscopy images of the fins reveal small round cells in the subsuperficial layer of the epidermis. (A-B, D-G) TEM images of a cross section through the epidermis of the fin. (A, B) The epidermis is composed of several layers of keratinocytes (Ker); the top layer being called pavement cells (PVC) with microplicae. Underneath the pavement cell layer, small round cells resemble HCS-cells. Bm: basement membrane. Col: collagen of the dermis. (A’, B’) Higher magnification of the putative HCS-cells. The cells contains a large nucleus (nuc) relative to its size, a Golgi apparatus (Ga), mitochondria (m) and vesicles (v). Unlike adjacent cells, the HCS-cells do not have desmosomes (d), however they are in contact with nerve fibers (n). (C) SEM image of the fin surface. The pavement cells (PVC) are covered with ridges called microplicae. (D-G) Higher magnification of features of HCS-cells in A’ and B’: Golgi apparatus (D), Nerve fiber adjacent to HCS cell and vesicle in the cytoplasm of HCS cell (E), large mitochondria (F), Neuronal projection touching HCS cells (G).

The EM imaging analysis revealed the presence of small roundish cells with a diameter of 2.87 ± 0.58 μm (n = 11), characterized by a high nucleus-to-cytoplasm ratio and a rather regular circumference (Fig. 4A, B). Their cytoplasm was polarized with more organelles towards one side, resembling the 5-HT/SV2 distribution in confocal analysis. The cells contained the Golgi apparatus, abundent mitochondria, and 50-100 nm wide secretory vesicles (Fig. 4A-B’, 4D-G). In accordance with immunofluorescence staining, the cells were not attached to the neighboring keratinocytes with desmosomes. At the periphery of the cells, we found nerve fibers (Fig 4E, G).

These cells did not contain any dense-core granules, the main feature of classical Merkel cells in tetrapods and fishes (Lane and Whitear, 1977; Tachibana et al., 1984; Zaccone, 1986; Halata et al., 2003). Furthermore, the small round cells did not bear an apical antenna-like protrusion, a key structure of chemosensory cells in zebrafish (Kotrschal et al., 1997). We concluded that the identified cells might represent a distinct type of paraneuronal cells in the fin. These cells could, however, be related to Merkel-like cells.

### Scattered distribution of HCS-paraneurons in the caudal fin

To investigate the distribution pattern of HCS-cells in the caudal fin, we performed immunofluorescence on whole mount specimens. We found that the fin surfaces contained numerous scattered 5-HT/SV2-positive cells (Fig. 5A). Consistent with the analysis of fin sections, a vast majority of 5-HT-labeled cells were also SV2-positive. However, in addition to the scattered SV2/serotonin-positive cells, we also found SV2-positive/serotonin-negative cell clusters, which were regularly distributed along a nerve fiber in parallel to rays (Fig. 5B). The chain-like arrangement of these cellular aggregates is reminiscent of neuromasts, which function as mechanoreceptors (Kniss et al., 2016). Consistent with previous studies (Dufourcq et al., 2006; Ghysen and Dambly-Chaudiere, 2007), we found that three to six rows of neuromasts were present in the caudal fin. By contrast, serotonin-positive individual cells were dispersed across the entire fin surface (Fig. 5A’-A’’).

**Figure 5.**
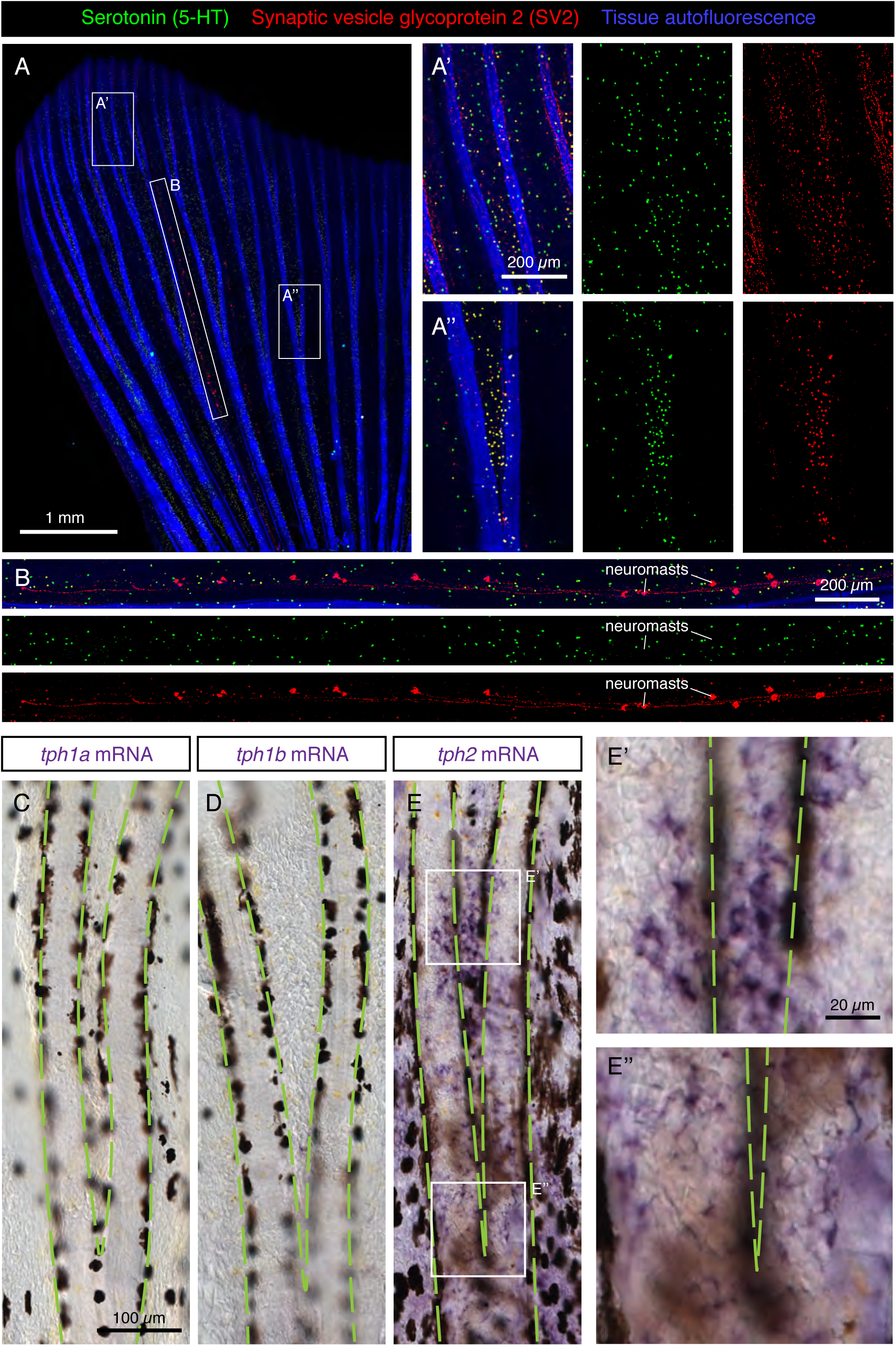
Scattered distribution of HCS-cells in uninjured adult fins. (A-B) Whole-mount immunofluorescence staining of uninjured fins for serotonin (5-HT; green) and SV2 (red). Tissue autofluorescence allows visualization of bony rays (blue). (A’-A’’) Higher magnifications of different fin regions. The area immediately above a ray bifurcation shows a higher density of HCS-cells. N = 12. (B) A zoom of the interray region marked by an elongated frame in (A). Aligned multicellular aggregates of SV2-positive and 5-HT-negative cells represent mechanosensory neuromasts. (C-E) Whole mount *in-situ* hybridization of uninjured fins for rate-limiting enzymes in the synthesis of serotonin *tph1a*, *tph1b* and *tph2*. Microscopic images near a bifurcation area (bifurcating bones highlighted with green dashed lines). *tph1a* and *tph1b* are not expressed, while *tph2* is expressed in a dotty pattern in the epidermis with a higher density between the bifurcating bones. N = 3 for each probe.

The serotonin synthesis pathway requires the conversion of the amino acid L-tryptophan to 5-hydroxytryptophan by the enzyme Tryptophan hydroxylase (Tph) (Berger et al., 2009). The product of this reaction is the precursor of serotonin. To determine whether this enzyme is expressed in the epidermis of the fin, we performed *in-situ* hybridization of whole fins with probes against the three homologous genes encoding it: *tph1a, tph1b and tph2*. Among them, we found a positive signal only for *tph2* (Fig. 5C-E). Closer magnification of the tissue revealed expression in single cells on the fin surface in a similar pattern to that of HCS-cells (Fig. 5E’-E’’). We concluded that the paraneuronal cells of the epidermis express *tph2*, to synthesize 5-HT.

To determine the density of HCS-cells in the caudal fin, we quantified the number of serotonin-positive cells per area of epidermis. We found that the densities of HCS-cells substantially varied even among sibling fish from 25 to 300 cells per mm^2^, with an average number of approx. 140 cells per mm^2^ (Fig. 6B). Given that the average surface of the analyzed fins was approx. 20 mm^2^ and that the fin has two epidermal surfaces, we calculated that the adult caudal fin contains roughly 6000 HCS-cells.

**Figure 6.**
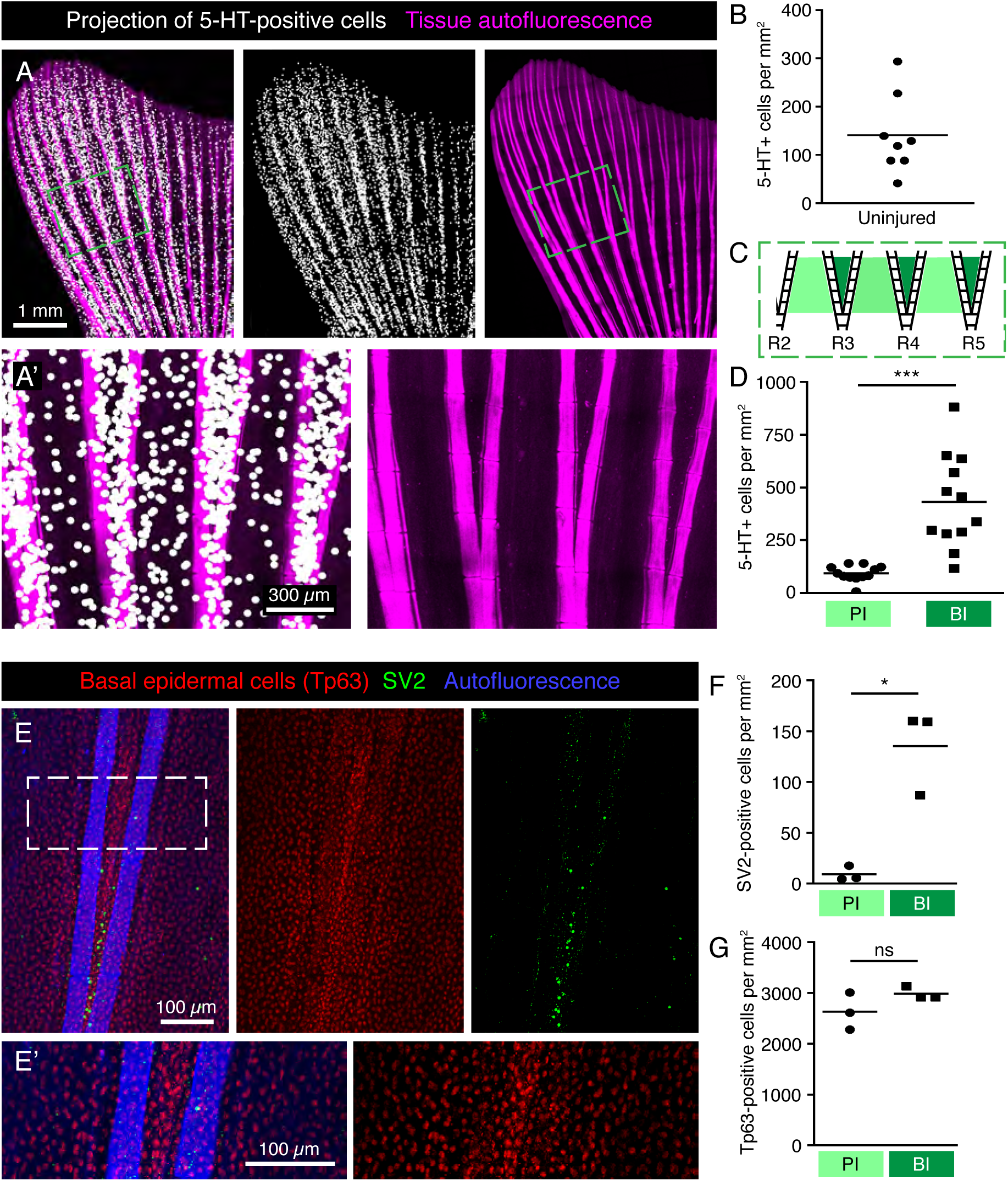
Specific pattern of HCS-cells around bifurcations in the uninjured fin. (A-A’) Projections of 5-HT-positive cells (white dots) onto fin surface with autofluorescent bones (pink) for better visualization of the HCS-cell pattern. One dot is projected at the position of each 5-HT-positive cell. (A’) Zoom of the framed area in C shows a higher density of cells between bifurcating rays. (B) Quantification of overall density of 5-HT-positive cells in uninjured fins shows high variability. Each dot represents one fin. The average density is 141 ± 82 cells per mm^2^. N = 8. (C) Schematic representation of an area near primary bifurcations used for the quantifications in (D). Primary interrays (light green) and the adjacent bifurcation interrays (dark green). (D) Quantification of 5-HT-positive cells in primary interray (PI) versus bifurcation interrays (BI) in a range of 5 ray segments after the bifurcation, as shown in (C). Density is approx. 4-times higher in bifurcation interrays (432 ± 222 vs 93 ± 38 cells per mm^2^). N = 12. ***: p < 0.001 (E-E’) Whole mount immunofluorescent staining for SV2 (green) and Tp63 (red), which strongly labels the nuclei of the epidermis. Keratinocytes do not display the same difference in the distribution pattern around bifurcations that is seen for HCS-cells. (F) Quantification of SV2-positive cells in PI versus BI in the bifurcation areas as outlined in (C). Density of SV2 cells is significantly higher in BI (9 ± 7 vs 135 ± 42 cells per mm^2^) *: p < 0.05 (G) Quantification of Tp63-positive cells in PI versus BI in the bifurcation areas. No significant difference is seen in Tp63 density between PI and BI (2630 ± 364 vs 2985 ± 125 cells per mm^2^). N = 3.

### Increased density of HCS-cells at the ray bifurcation areas

To increase the visibility of serotonin-positive cells on images of the whole fin surfaces, we performed a computational projection of these cells as enlarged dots. We anticipated that this method could highlight any local differences of cell density in the tissue. Indeed, these projections revealed that the distribution of HCS-cells was uneven (Fig. 6A). First, the density of HCS-cells was different between the lateral lobes and the middle part of the fin. Specifically, the fin region supported by the 6 lateral rays contained a twice-higher HCS-cells density as compared to the fin region spanned by the 4 central rays (lateral fin part: 165 ± 80 cells per mm^2^ versus medial fin part: 68 ± 48, p<0.001). Second, some local differences were visible around the ray bifurcations (Fig. 6A’, C, D). Namely, the number of HCS-cells was 4-fold higher in the interrays above the bifurcation point (bifurcation interray, BI), as compared to the neighboring primary interrays adjacent to the rays (primary interray, PI) (Fig. 6D).

The architecture of ray bifurcation might also be associated with changes in density of keratinocytes. To test this and quantify keratinocytes in the relevant areas of the fin, we used the nuclear marker Tp63 (tumor protein 63, p63), which is expressed in basal epidermal cells of the zebrafish fin (Stewart and Stankunas, 2012). The densities of Tp63-positive nuclei in the bifurcation and primary interrays were similar, suggesting no difference in the density of keratinocytes around bifurcations (Fig. 6E-G). We concluded that the increased density of HCS-cells at the bifurcation area is not caused by local changes in epidermal density.

The vital fluorescent dye DASPEI has been previously used as a marker of hair cells of neuromasts, electroreceptors and chloride cells in the teleost skin (Foskett et al., 1981; Jørgensen, 1992; Whitfield et al., 1996; Dufourcq et al., 2006; Froehlicher et al., 2009). The cellular entry of this dye is thought to depend on the expression of specific types of ion channels and carriers in the plasma membrane. To test whether HCS-cells absorb DASPEI, we incubated fish in water containing this compound and performed live-imaging of fins. Consistent with previous reports (Dufourcq et al., 2006; Froehlicher et al., 2009), the neuromasts were demarcated by the fluorescent dye (Fig. 7B-C). In addition, we found some scattered DASPEI-labeled cells. However, the DASPEI-positive cells displayed a different distribution pattern from HCS-cells (Fig. 7A-C). Indeed, the density of DASPEI-positive cells was overall lower than that of HCS-cells (Fic. 7D) and the pattern around the bifurcation areas was not observed (Fig. 7E). This finding indicates that HCS-cells did not absorb DASPEI, suggesting that these cells do not reach the surface of the epithelium or cannot absorb this compound.

**Figure 7.**
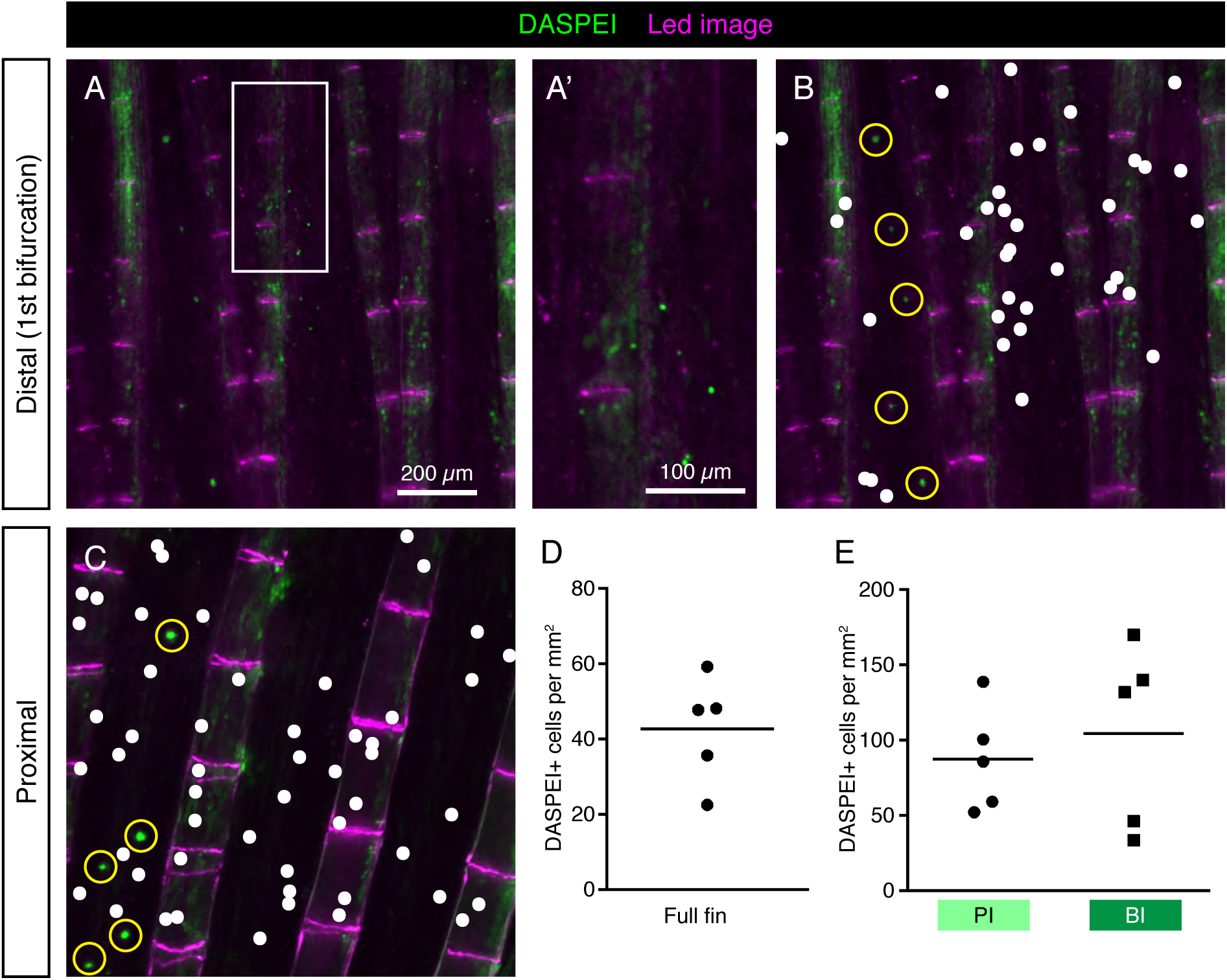
DASPEI-positive cells in the uninjured fin do not display a HCS-like distribution pattern. (A-C) Live imaging of DASPEI-stained (green) uninjured fins. (A-A’) Image of a fin around the first bifurcation level. No specific pattern of DASPEI-positive cells is observed around the bifurcation. (B) The same image as shown in (A) with a white dot projections of DASPEI-positive cells. Larger DASPEI-positive bundels aligned between the rays are neuromasts (yellow circles). (C) Image of a fin in the proximal region of the fin with white dots projected over solitary DASPEI-positive cells. DASPEI-positive neuromasts are clearly visible (yellow circle). (D) Quantification of overall density of DASPEI-positive solitary cells per mm^2^ in the fins. Density Overall density was 42 ± 15 cells per mm^2^, much lower that the overall density of HCS-cells. (E) Density pattern around bifurcation areas quantified according to the areas highlighted in Fig 6C. Density of DASPEI-positive cells in the PI was not significantly different from the density in th BI (87 ± 35 vs 104 ± 61 cells per mm^2^). N = 5

To determine whether the distribution pattern of HCS-cells is robust, we analyzed fins of *another longfin* (*alf*) fish, which have a gain-of-function mutation in the potassium channel gene *kcnk5b* and display severely elongated fins (Perathoner et al., 2014). The *alf* fins showed a similar pattern of HCS-cells along the lateral-medial axis (Fig. 8A), although the overall density of HCS-cells was higher in *alf* fins compared to wild-type fins (Fig. 8B). Furthermore, the areas of bifurcations were associated with increased number of HCS-cells as compared to the adjacent primary interrays, like in wild type fish (Fig. 8A’, C). Overall, the distribution of HCS-cells in the fin is not random, but seems to be regulated by positional cues of the appendage, even in morphologically elongated fins.

**Figure 8.**
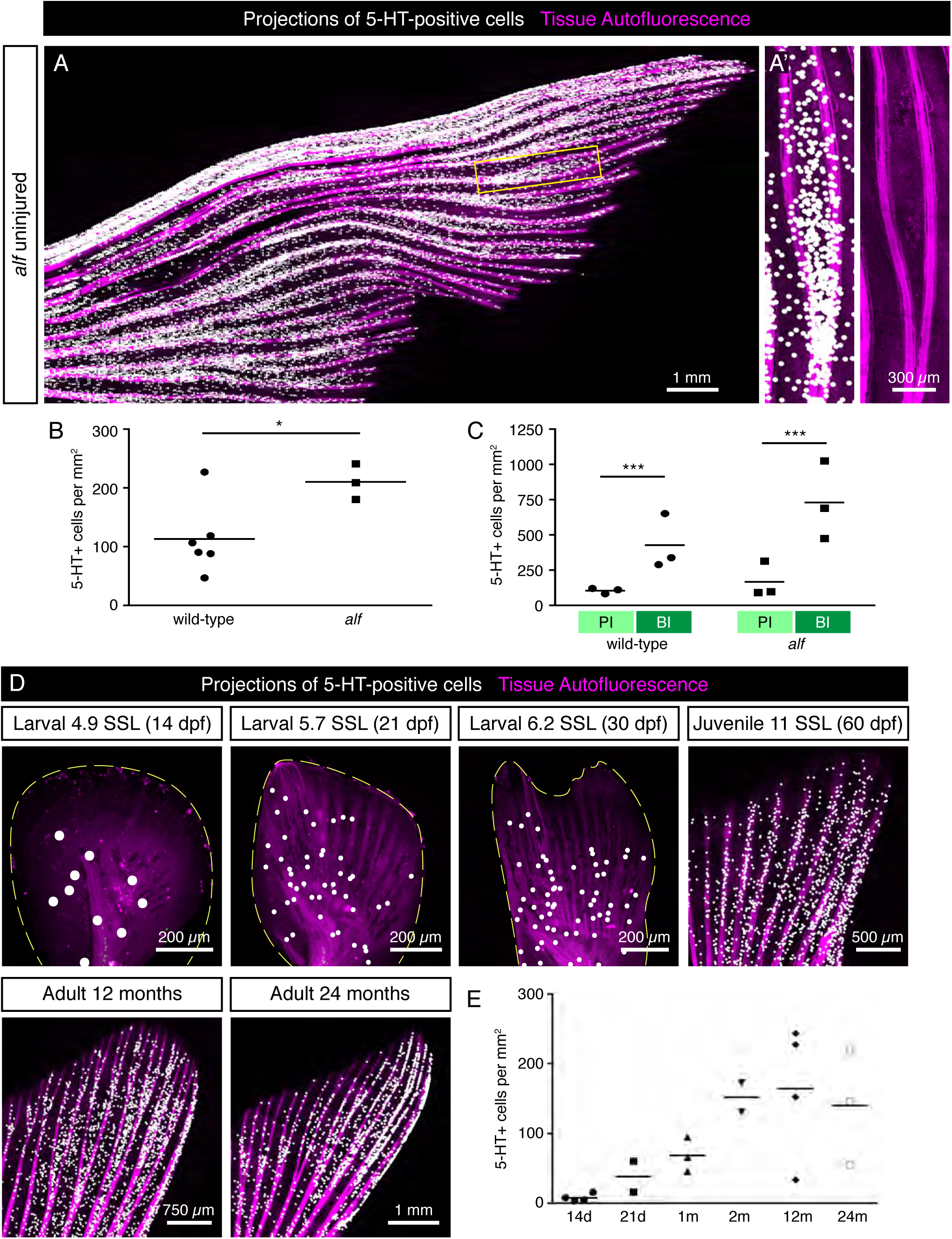
Distribution of HCS-cells in *alf* mutants and during ontogenesis of wild type zebrafish. (A) Projections of 5-HT-positive cells (white dots) in an uninjured *alf* fin based on immunofluorescence staining of the rays. Autofluorescence of tissue with bones (pink). (A’) Zoom of the framed region in (A) shows higher density of HCS-cells in bifurcation interrays, like in wild type fins. (B) Quantification of HCS-cell density in *alf* fins versus wild-type fins shows a significantly higher overall density of HCS-cells in *alf* fins (113 ± 60 vs 210 ± 30 cells per mm^2^). N ≥ 3. *: p < 0.05 (C) Quantification of HCS-cell density in primary interray (PI) vs bifurcation interray (BI) in wild-type versus *alf* fins. Density difference is present in *alf* fins (104 ± 20 vs 426 ± 196 cells per mm^2^ in wild type and 166 ± 112 vs 729 ± 277 in *alf*). N ≥ 3. ***: p < 0.001 (D) Projections of 5-HT-positive cells in fins at different ages ranging from 14 days to 24 months post-fertilization, based on immunofluorescence staining. For juvenile stages, the developmental stage is shown as standardized standard length (Parichy et al., 2009). (E) Quantification of density of 5-HT-positive cells in caudal fins during post-embryonic ontogenesis. While the density is lower at larval stages, once fins switch to a bi-lobed morphology, the concentration of HCS-cells increases and remains constant through the adult life of the fish (7 ± 5 at 14 dpf; 38 ± 30 at 21 dpf; 68 ± 25 at 30 dpf; 152 ± 60 at 60 dpf; 164 ± 95 at 12 months and 140 ± 82 at 24 months). N ≥ 2.

The zebrafish caudal fin undergoes morphological changes during post-embryonic development (Goldsmith et al., 2006; Parichy et al., 2009). To determine whether the number of HCS-cells changes during the lifespan, we performed analysis of 5-HT-positive cells of caudal fins at different ages starting at the post-embryonic stage. For morphological staging, we classified fish according to their standardized standard length (SSL) (Parichy et al., 2009). At SSL 5, corresponding to 14 dpf, before metamorphosis of the fin, the density of 5-HT-positive cells was very low around 10 cells/mm^2^ (Fig. 8D-E). During metamorphosis at SSL 5.7 to 6.2 (21 dpf to 30 dpf), as the fin switches from a paddle-shape to a bilobal shape, the density of 5-HT-positive cells increased, concomitant with the formation of rays. Finally, the 5-HT-positive cells reached the adult density of about 140 cells/mm^2^ around 60 dpf (SSL 12) and remained unchanged through the adult life of the fish (Fig. 8D-E). Thus, the high density of HCS-cells is established during metamorphosis and remains largely constant during adult life.

### Fluid dynamics in a model of a ray bifurcation correlate with the HCS-cell distribution

Examination of the HCS-cells distribution in the caudal fin revealed a higher density of these cells inside the bifurcation interrays, as compared to the adjacent primary interrays. We hypothesized that specific fin structures may be exposed to different flow velocities, whereby the accumulation of HCS-cells occurs at the regions that are exposed to optimal conditions for water sampling. To test this idea, we constructed models of a fin using a plate with rods mimicking bones with and without bifurcation, taking into account various physical parameters (Fig. 9, Fig. 10A, B). The models were placed inside a flow chamber in such a manner that swimming motion could be imitated through oscillation of the plates. The fluid velocities were analyzed based on high-speed recordings of beads flowing in the water with 3 cameras, allowing for the analysis of flow in 3 dimensions. Average values of the parallel (streamwise) and perpendicular components of the fluid velocity were analyzed for the bifurcated interray region (BI) and the adjacent primary interray regions (PI). These models brought to light several flow features characteristic to the zone located inside the V-shape bifurcation.

**Figure 9.**
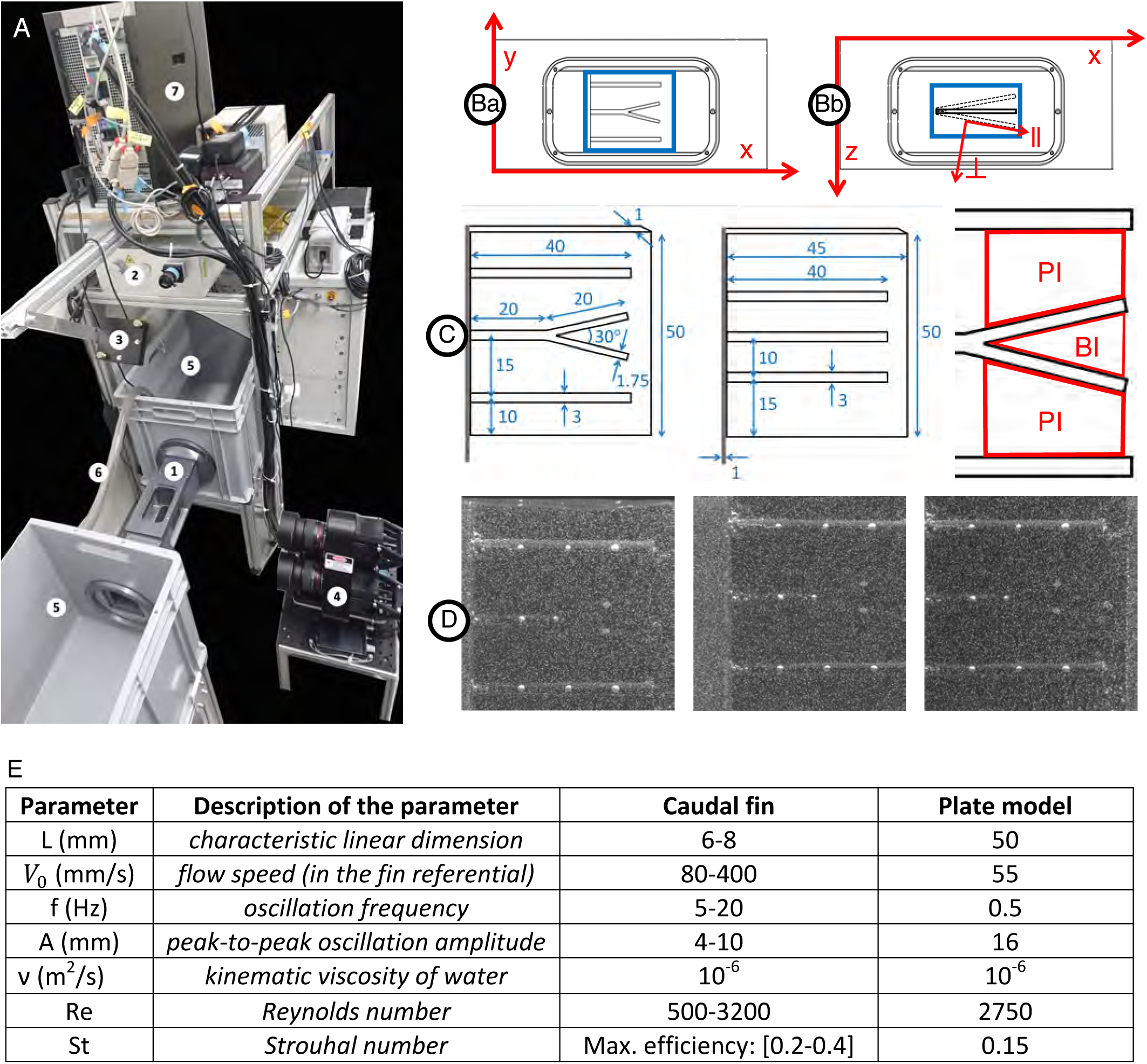
Experimental setup for hydrodynamics measurement. (A1) Flow chamber with transparent windows on three sides and a fixation wall on one side for inserting the fin model actuated with a servomotor fixed outside the chamber. (Ba) Close-up of the flow chamber (frontal view with x and y axes) with interrogation volume in blue (∼50×50 mm^L^) used for the 3D reconstruction of particles positions. (Bb) Close-up of the flow chamber (top view with x and z axes) with interrogation volume in blue (∼50×20 mm^L^) used for the 3D reconstruction of particles positions. The oscillation of the plate model is illustrated including the parallel (║) and perpendicular (⊥) axes moving with the plate and used for the velocity vector decomposition. (A2) 200 mJ dual-head pulsed Nd:YAG laser equipped with a pair of cylindrical lenses to expand the beam. (A3) Mirror to deflect the laser beam and illuminate the volume inside the flow chamber. (A4) Three cameras (4 MP, 85 mm lenses) mounted on a plate in a triangular arrangement, pointing at the flow chamber to image the 3D flow based on the triangulation principle. The distance between the cameras plate and the center of the water tunnel is ∼46.5 cm. (A5) Water tanks connected to both sides of the flow chamber in a recirculating system. (A6) Pipe connected to a pump carrying water from one tank to the other to control the flow inside the chamber. (A7) V3V software and synchronizer to control the timing of the laser pulses and the opening of the camera apertures. (C) Models of a fin consisting of a rigid plate supporting half-cylindrical rods including a centered bifurcated ray (left), control plate model with straight rods only (middle) and sketch of the primary (PI) and bifurcation (BI) interrays. All dimensions indicated in mm. (D) Example of raw triplet images (captured by the left, right and top cameras respectively) with the illuminated tracer particles (∼50 µm diameter) and the fin plate model rendered visible by the addition of equally spaced white dots painted directly on the rods, allowing for surface tracking throughout the oscillation period. (E) Parameters of the plate model as compared to a fin.

**Figure 10.**
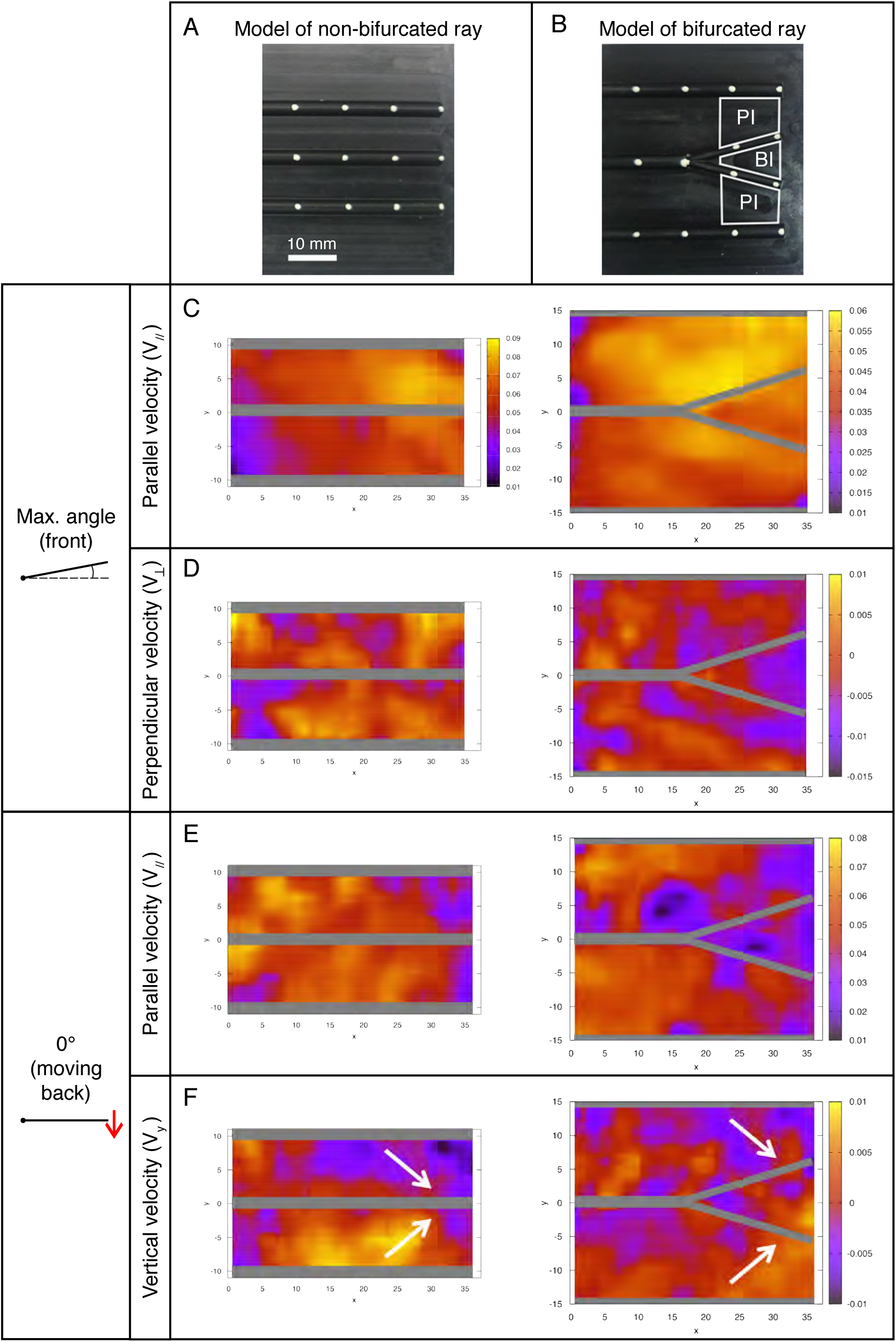
Fluid velocity profiles in the bifurcation interray zone (BI) and the primary interray zone (PI) highlight particular fluid motion at the interray bifurcation site. (A, B) Models of a fin consisting of a rigid plate supporting half-cylindrical rods used for the hydrodynamic profiles study in a version with three parallel rods (A), and a bifurcation in the central ray (B). The indicated primary interray areas (PI) and bifurcation interray area (BI) are used for the velocity components averages presented below. (C-F) Flow profiles (all color scales in meters per second) at two key positions during the plate’s oscillation period. The corresponding average measurements are shown in (Fig. 6B and C). (C, E) A lower fluid velocity parallel to the surface (V_//_) is typically associated with a thicker boundary layer, and hence a potentially better detection of chemical signal in the bifurcation region, owing to longer time reaction for the cellular receptors and better averaging of the signal. (D) Larger absolute values of the perpendicular velocity (V⏊< 0) indicate a higher rate of fluid motion directly towards the bifurcation region and imply a better access to molecules advected by the fluid from outside the boundary layer. (F) The vertical velocity component (V_y_) also reveals a global fluid motion towards the inter-bifurcation region, naturally correlated with a movement perpendicularly away from the fin as indicated by the positive value of V⏊ in (Fig. 6C). The velocity fields correspond to averages over three (0° angle) to five timeframes (max. angle).

Key positions were selected along a full period of oscillation, namely the starting position (0°) of the forward and backward movements (Fig. 11A, C), followed by the end-position in both directions (Fig. 11B, D).

**Figure 11.**
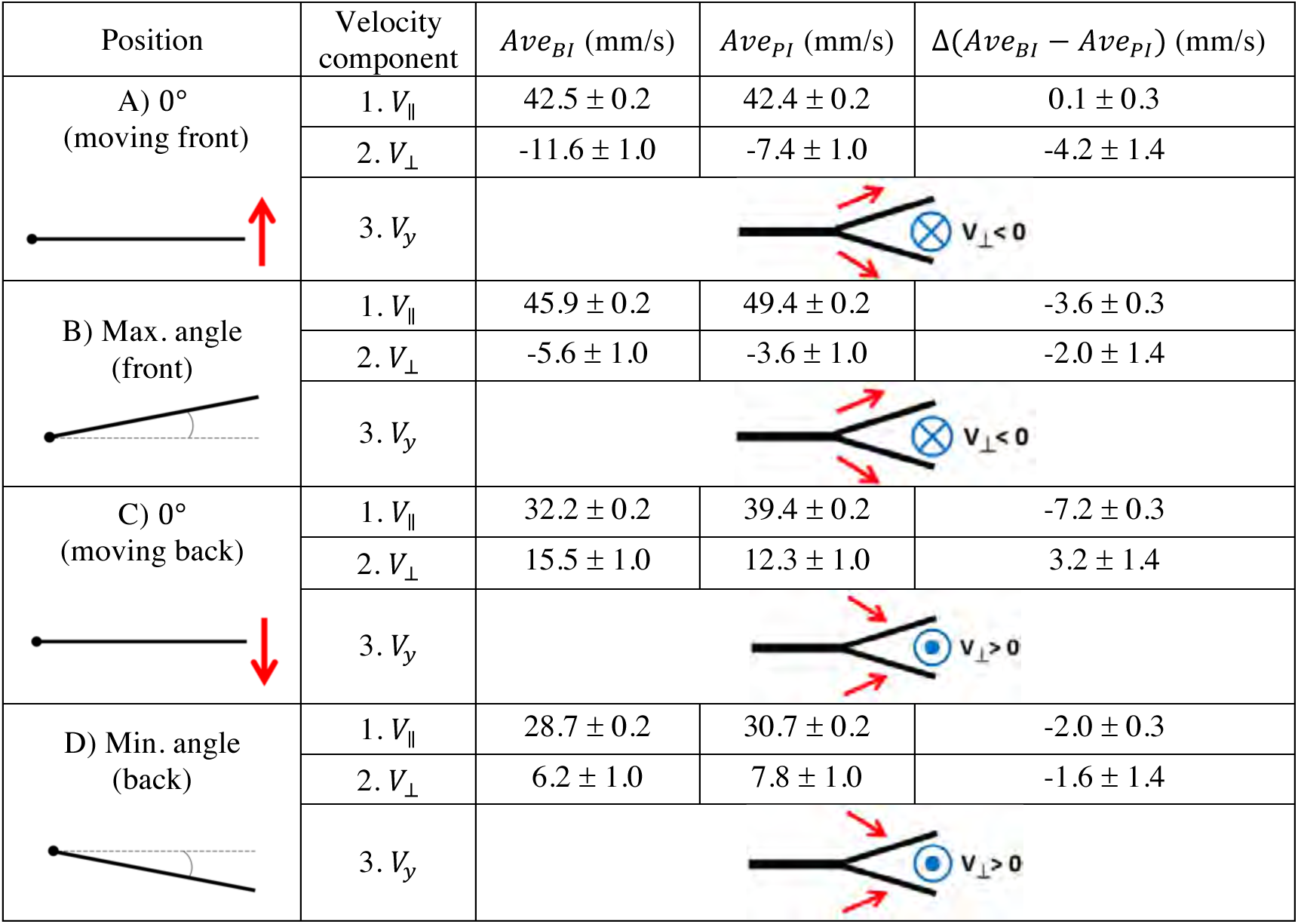
Quantification of fluid profiles in the bifurcation interray zone (BI) and the primary interray zone (PI), at four equidistant positions of the fin model during a complete period. (A-D) Four key points were selected along a full period of oscillation as indicated in the left column. A schematic depiction of the vertical velocity component (V_y_) is represented by the simplified diagrams in the right column, in which the red arrows symbolize the V_y_ vectors, pointing towards or away from the bifurcation region. The blue symbols represent the resulting fluid motion in the perpendicular direction, namely the V_⊥_ vectors, pointing outward (circle with a dot in the middle) or inward (circle with a cross in the middle). The average velocity components parallel and perpendicular to the plate (V_//_ and V_⊥_) are indicated in millimeters per second. They are noted Ave_BI_ or Ave_PI_ depending on the averaging surface (BI region or PI region), and the difference between the averages Δ(Ave_BI_ - Ave_PI_) is also shown.

We considered the implications of a hypothetical mechanosensory function not observed yet on fish fins, more specifically, the capacity to encode local flow signals. The thickness of the boundary layer, the layer of fluid which adheres to the body surface resulting in slower fluid motion, over the plate model was measured at three motion instants: a stationary reference case, where the plate was kept immobile, as well as the 0° angle position during the forward and backward motion of the plate. These results are illustrated in Fig. 12A, where each panel presents the boundary layer thickness over the two versions of the plate model: with the straight rods and with the central bifurcated ray. For the latter geometry, the boundary layer was measured at two locations on the plate, corresponding to the primary interray and the bifurcation interray. The boundary layer thickness of the stationary reference model, without bifurcations, appears to be relatively constant along the plate, oscillating in the range between 2 mm and 3.5 mm until the tip (top panel, triangles). For the straight rods, the most striking effect induced by the pitching motion was an increase of the boundary layer thickness during forward motion, except at the very distal tip, where the boundary becomes much thinner (middle panel, triangles). A qualitatively inverse phenomenon is observed during backward motion, where the boundary layer thickness increases everywhere except at the most upstream, proximal position, where it is thinner (bottom panel, triangles). For the stationary bifurcated rod model, the boundary layer thickness in the primary interray is larger than that of the parallel rods geometry, with a value situated between 5 mm and 6 mm approximately, except at the most proximal portion of the plate (*x* < 5 mm), where the reference velocity is not reached, which shows on the graph as an infinitely large boundary layer (top panel, stars). The pitching motion has a similar influence in the primary interray of the bifurcated ray as in the parallel rods geometry: the thickness is reduced at the distal tip during forward motion, and at the proximal portion during backward motion (stars in the middle and bottom panels respectively). In each motion, the points corresponding to the boundary layer thickness inside the bifurcation interray roughly overlap with the thickness in the primary interray (circles in top, middle and bottom panels). However, the numerous missing points imply that the average velocity is not reached at those *x* positions, a sign that distinct flow profiles emerge in both regions of the plate (PI versus BI). The different flow fields generated in the primary and bifurcation interrays are illustrated in Fig. 12B. For the three motion instants (immobile, 0° moving forward and 0° moving backward), top views show the normalized streamwise fluid velocity (*V*_║_/*V*_║’()_) in cross-sections of the measurement volume located at the level of the PI and BI regions. In all three cases, the presence of the bifurcation (indicated by a white bar) clearly disturbs the flow.

**Figure 12.**
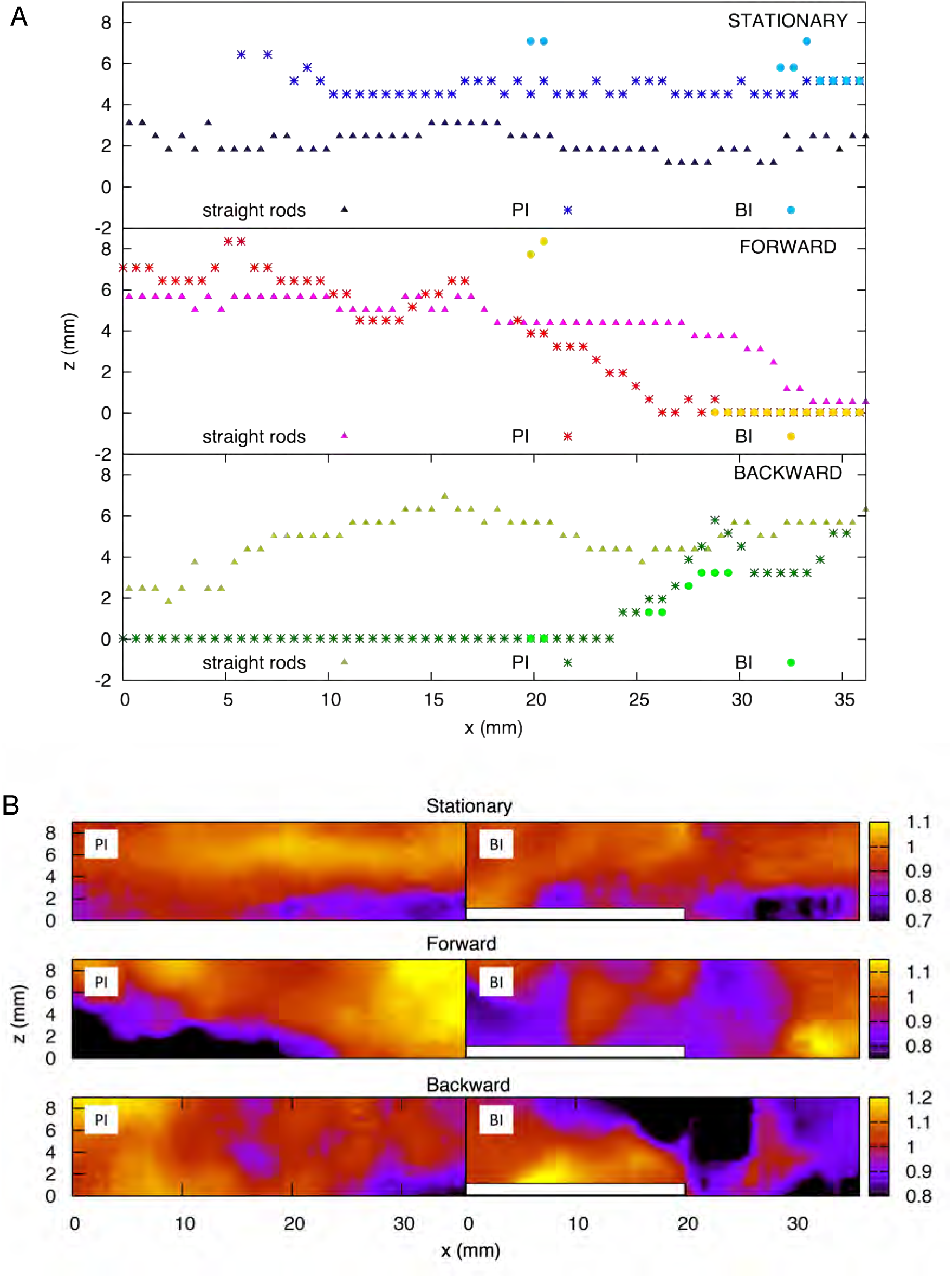
Analysis of the boundary layer on scaled fin models. (A) Boundary layer thickness for the plate model with straight rods, and for the plate model with a bifurcated middle rod, at the primary interray and the bifurcation interray zones. Top panel: Stationary plate. Middle panel: Zero-degree angle position during forward motion of the plate. Bottom panel: Zero-degree angle position during backward motion of the plate. (B) Color map of the streamwise velocity component (*V*_║_) normalized by the average value - top view of the flow chamber vis-a-vis the primary interray (PI) and the bifurcation interray (BI). Top panel: Stationary plate. Middle panel: Zero-degree angle position during forward motion of the plate. Bottom panel: Zero-degree angle position during backward motion of the plate.

The flow at each of the selected positions of the plate is also represented in Fig. 10, where a frontal view of the bifurcation model is shown. The different flow patterns observed in that figure are summarized in Fig. 11, using simplified schematic diagrams which globally illustrate the vertical motion of the fluid with respect to the bifurcation point (*V*_y_, symbolized with red arrows), together with the resulting perpendicular fluid motion at that same point (*V*_⊥_, noted with a blue symbolic vector pointing in or out of the plane). The observation encapsulated by these diagrams is that during the forward motion and until the plate has reached maximum angle, the vertical fluid motion is globally directed away from the bifurcation interray, resulting in a perpendicular fluid motion pointing towards BI (Fig. 11A, B). Moreover, in both positions, the average value of *V*_⊥_, is larger (in absolute value) in the bifurcation interray versus the primary interray. This phenomenon is also visible in flow profiles depicted in a color map (Fig. 10D). No such increased flow towards the plate is observed in the plate lacking a bifurcation.

By contrast, during the backward portion of the oscillation, the fluid flows vertically towards the bifurcation zone and as a consequence it streams perpendicularly away from that area (Fig. 11C, D and Fig. 10F). An additional striking effect of the ray bifurcation on flow is observed in three of the selected angular positions (0° moving back, and at minimum and maximum angle), where the parallel component of the flow is reduced inside the BI area in comparison to the PI area (Fig. 11B-D). Accordingly, averages of *V*_║_ are lower in the bifurcation zone for these time points, although the flow remains globally in the upstream direction (Fig. 11B-D). This effect is also visually represented in flow profiles (Fig. 10C, E).

The observed flow velocities can be interpreted as a favoring factor for the enhanced presence of HCS-cells inside the V-shape interray. Lower fluid velocity parallel to the surface inside the bifurcation region is indicative of a thicker boundary layer, namely the layer of fluid which adheres to the body surface resulting in slower fluid motion relative to the object. Moreover, the presence of the bifurcated ray in the flow is likely to induce a transition to turbulence. Although the exact nature of the hydrodynamic disturbances induced in the bifurcation interray requires further investigation, the data constitute evidence that preferential flow conditions are present in that region for potential sensory cells.

### Efficient restoration of HCS-cells during fin regeneration

After an amputation, zebrafish fins can fully regenerate within approx. 3 to 4 weeks (Pfefferli and Jaźwińska, 2015; Wehner and Weidinger, 2015). To determine whether the population of HCS-cells is restored during fin regeneration, we performed whole mount immunofluorescence analysis at different time points after amputation. Analysis of fins shortly after blastema formation at 2 days post-amputation (dpa) revealed re-appearance of 5-HT-positive cells in the new tissue that protruded distal to the amputation plane (Fig. 13B). At the subsequent regenerative stages, at 7, 14 and 30 dpa, the new outgrowth comprised HCS-cells in similar numbers and pattern as in uninjured fins (Fig. 13A-F). Quantification of the cells revealed that the density of HCS-cells ranged from 100 to 150 cells per mm^2^ at all regenerative stages (no significant differences between regenerative time points and uninjured fins) (Fig. 13F). Analysis of bifurcations showed reappearance of the uneven density between bifurcation interrays and primary interrays at 30 dpa (Fig. 13E’, G). This pattern could already be observed at 14 dpa if the bifurcations were already present (Fig. 13D’). Taken together, HCS-cells and their specific pattern can be efficiently reestablished during fin regeneration.

**Figure 13.**
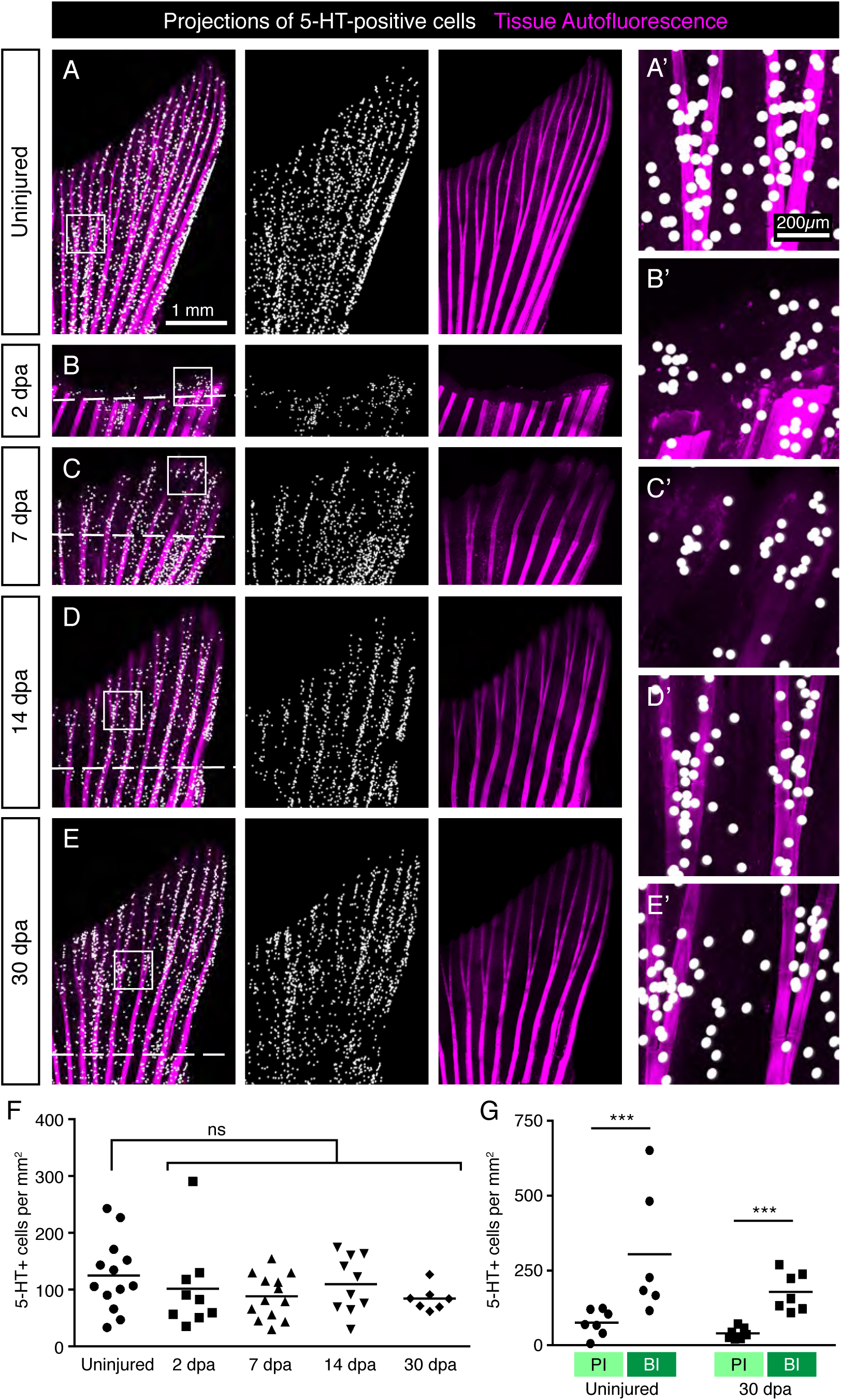
Restoration of HCS-cells during fin regeneration after amputation. (A-E) Projections of 5-HT-positive cells (as detected by immunofluorescence) on whole mount uninjured fins, at 2 days post-amputation (dpa), 7 dpa, 14 dpa and 30 dpa. The formation of the new outgrowth distal to the amputation plane (dashed line) is associated with concomitant restoration of HCS-cells. (F) Quantification of overall HCS-cell density at the indicated time points. No significant (ns) difference in density during regeneration compared with uninjured fins (126 ± 67 cells per mm^2^ in uninjured; 102 ± 77 at 2 dpa; 88 ± 39 at 7 dpa; 109 ± 50 at 14 dpa; 84 ± 21 at 30 dpa). N ≥ 7. (G) Quantification of HCS-cell density in primary interray (PI) vs bifurcation interray (BI) in uninjured fins vs fin regenerates at 30 dpa, as illustrated in (Fig. 6C). Differential density is reestablished after regeneration (75 ± 43 vs 386 ± 291 cells per mm^2^ in uninjured and 40 ± 18 vs 178 ± 64 at 30 dpa). N ≥ 7. ***: p < 0.001.

To test, whether differentiated HCS-cells possess a mitotic activity, we used an antibody against Proliferating Cell Nuclear Antigen (PCNA), which is a marker of the G1/S phase of the cell cycle. We performed immunolabeling with SV2 and PCNA on sagittal sections through the epidermis. We found that very few SV2/PCNA-double positive cells were present on the fin surface. Indeed only 2.3% of SV2 cells were PCNA positive, while 14% of the surrounding epidermal cells were PCNA positive (Fig. 14A, B). This indicates that mature HCS-cells likely have a very low proliferative activity.

**Figure 14.**
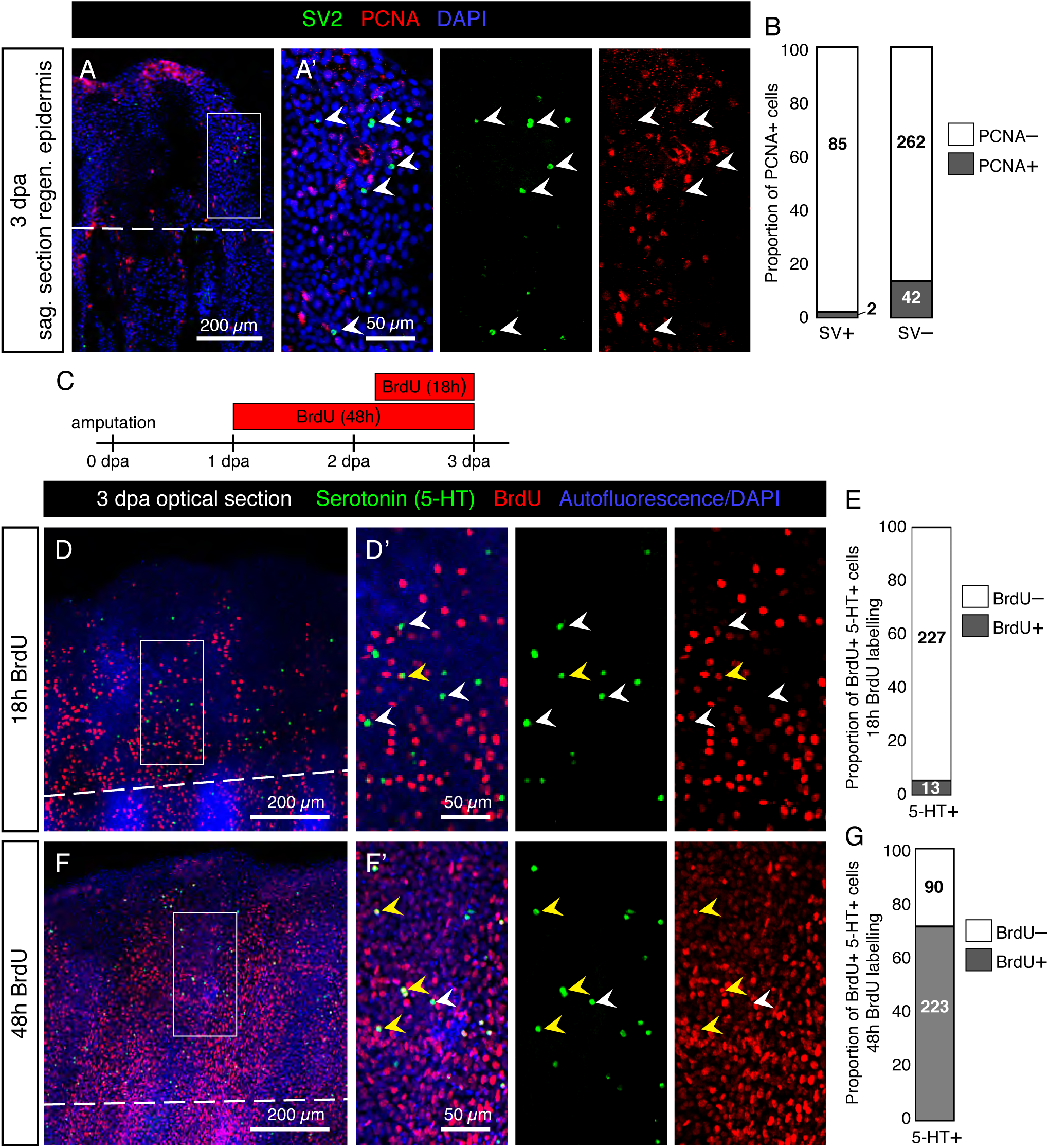
Low proliferative rate of mature HCS-cells during fin regeneration. (A) Immunofluorescent staining for SV2 (green) and PCNA (red) on sagittal section of fin wound epidermis at 3 dpa. (A’) SV2-positive cells are PCNA-negative (arrowheads). (B) Proportion of PCNA-positive cells among SV2-positive HCS-cells versus in SV2-negative epidermal cells (2.3% PCNA-positive in HCS-cells versus 13.8% PCNA-positive in other cells). N = 3. Number of cells for each group indicated on graph. (C) Experimental design for BrdU assays. (D, F) Immunofluorescence staining of whole mount fins at 3 dpa for serotonin (green) and BrdU (red) after 18h or 48h of BrdU-treatment. The image shows confocal imaging through the epidermis distal to the amputation plane (dashed line). (D’, F’) Zoomed areas show BrdU-negative (white arrowhead) and BrdU-positive (yellow arrowhead) 5-HT-positive cells. (E, G) Proportion of BrdU-positive cells among 5-HT-positive cells after 18h and 48h of labeling (5.4% and 71%, respectively). N = 3 fins; total number of cells in each group indicated on graph.

To determine whether serotonin-positive HCS-cells derive from cells which proliferated after amputation and how long the differentiation process takes, we performed a BrdU labeling assay. We collected fins at 3 dpa after 18h or 48h of BrdU treatment (Fig. 14C). After 18h of BrdU labeling, immunofluorescence analysis of BrdU and 5-HT on whole mount fins revealed only very few double positive cells in the epidermis (Fig. 14D-E). Confocal imaging of the fin surface displayed predominantly 5-HT-positive/BrdU-negative cells, while adjacent epidermal cells were abundantly labelled by BrdU immunostaining. However, after 48h of BrdU treatment, a majority of 5-HT-positive cells were also BrdU-positive (Fig. 14F-G). This indicates that new HCS-cells are generated in the regenerating fin, and the differentiation process from the time of BrdU incorporation to the production of serotonin takes between 18 and 48 hours for most cells. Based on both assays, we concluded that differentiated HCS-cells are rather non-proliferative cells in the fin regeneration context, and probably originate from the proliferation of another cell type, followed by a differentiation process.

### Regeneration of HCS-cells is not affected by the inhibition of serotonin biosynthesis

The synthesis pathway of serotonin is dependent on Trypophan hydroxylase (Tph) (Berger et al., 2009). To test whether 5-HT is required for regeneration of HCS-cells, we applied a chemical Tph-antagonist, para-chlorophenylalanine (pcpa) (Jequier et al., 1967), which was previously validated in analytical and behavioral assays in zebrafish embryos and adults (Njagi et al., 2010; Airhart et al., 2012). Consistent with our previous study (König and Jazwinska, 2019), we found that treatment with 5 mM pcpa, starting from one day before fin clipping until 3 days post-amputation, did not affect fin regeneration (Fig. 15A-D). This treatment efficiently abolished 5-HT-immunoreactivity in HCS-cells, as examined on whole mount immunofluorescent stainings at 3 dpa (Fig. 15A, B, E). Despite the absence of 5-HT, HCS-cells could be detected by Calretinin/SV2-labeling and their density was unaffected by the inhibition of serotonin synthesis (Fig. 15C-D, F-G). We concluded that, while serotonin is a hallmark of this cell population, its production is not required for the generation of new HCS-cells in the fin regenerates.

**Figure 15.**
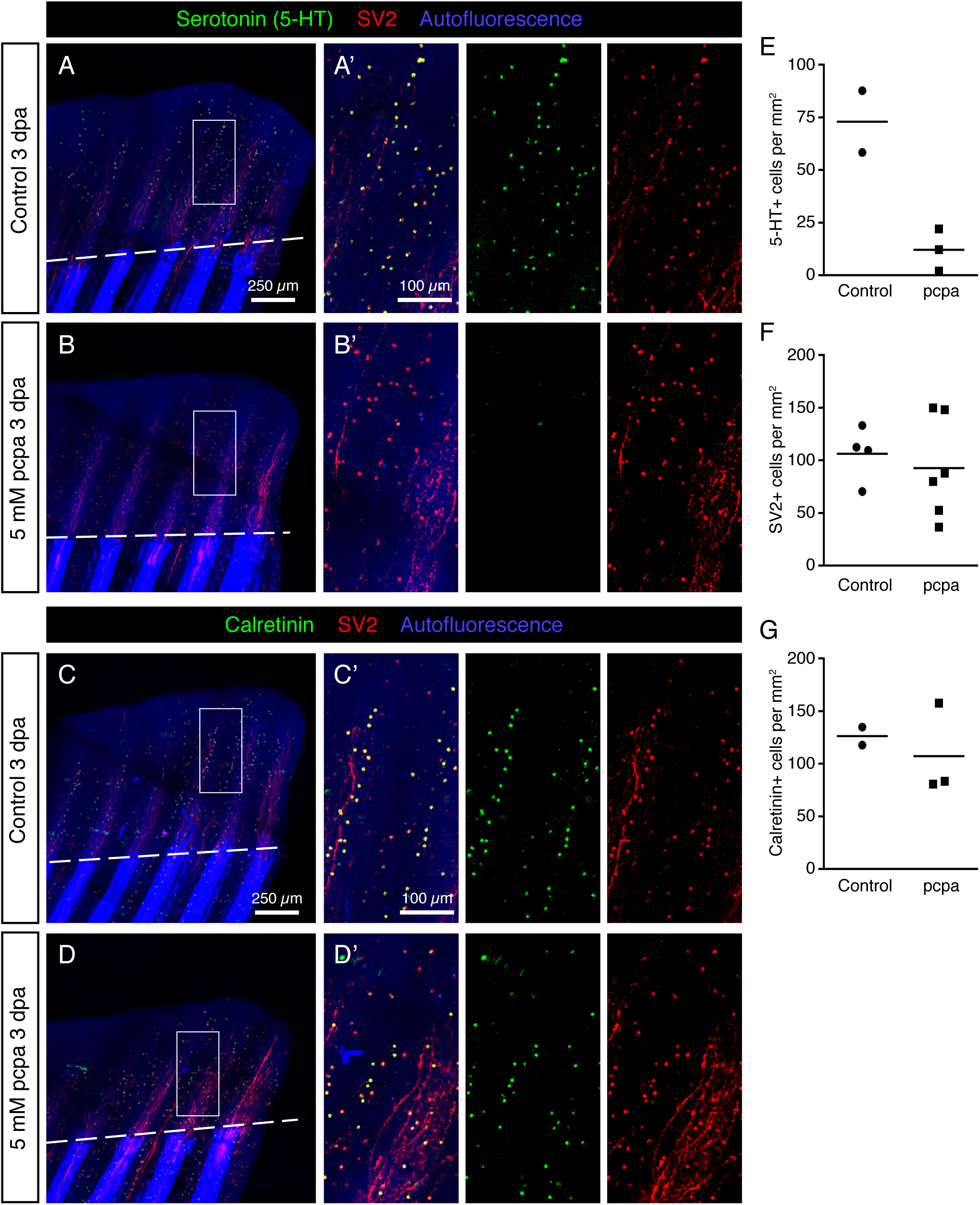
Inhibition of Serotonin production by pcpa-treatment does not prevent HCS-cell regeneration. (A-B) Whole mount immunofluorescent staining for 5-HT and SV2 in pcpa-treated (B) and control (A) fins at 3 dpa. Pcpa treatment eliminates serotonin, but does not eliminate SV2 expression in HCS-cells (A’-B’) The treatment does not prevent HCS-cell regeneration. (C-D) Whole mount immunofluorescence staining for Calretinin and SV2 in pcpa-treated (D) and control (C) fins at 3 dpa. Pcpa-treatment does not suppress the Calretinin expression in SV2-positive cells (C’-D’). (E) Density of 5-HT-positive cells in pcpa-treated fins decreased dramatically as compared to control fins, validating the activity of pcpa (72 ± 21 vs 12 ± 10 cells per mm^2^). N ≥ 3. (F) Density of SV2-positive cells is similar in control and pcpa-treated fins (106 ± 26 vs 92 ± 47 cell per mm^2^). N ≥ 3. (G) Density of Calretinin-positive cells remains unaltered despite of inhibition of serotonin production by the pcpa-treatment (126 ± 12 vs 107 ± 44 cells per mm^2^). N ≥ 3.

## Discussion

In adult zebrafish, the caudal fin propels the fish during swimming. The surface of this locomotory appendage is exposed to water flows which carry mechanical and chemical information. Thus, the epidermis of the fin may play a role in the focal detection of environmental signals. The recognition of stimuli and the translation of the received information occurs through sensors of the cutaneous neuroendocrine system. In this study, we investigated the distribution and regeneration of a specific class of serotoninergic paraneurons in the fin epidermis, which might participate in the detection and transmission of environmental cues in adult zebrafish.

### Molecular markers of solitary paraneurons in the epidermis of the caudal fin

A fundamental feature of paraneuronal cells is their regulated secretory pathway. This mechanism depends on storage vesicles with messenger substances which are released through calcium-dependent exocytosis in response to signals (Moghadam and Jackson, 2013). Synaptic and endocrine secretory vesicles possess a common transmembrane glycoprotein which is recognized by an anti-SV2 monoclonal antibody (Buckley and Kelly, 1985). Our study revealed that this antibody labeled two types of cells in the caudal fin, namely mechanoreceptive neuromasts and solitary dispersed cells. While neuromasts have been previously characterized in adult zebrafish (Dufourcq et al., 2006; Ghysen and Dambly-Chaudiere, 2007), the solitary cells we observed had not yet been investigated, to our knowledge. We found that these solitary cells, as opposed to neuromasts, were positive for three different serotonin antibodies. High-resolution confocal imaging showed a colocalization between 5-HT and SV2, suggesting that the storage vesicles of these cells contain this neurotransmitter. Furthermore, these cells contained Calretinin, which is a Ca^2+^ signaling dependent regulator of exocytosis and may regulate sensory functions (Schwaller, 2014; Pangršič et al., 2015; Soulika et al., 2016). The coexpression of all three markers, 5-HT, Calretinin and SV2 (HCS, a combination of the first letters of the markers), strongly suggests the paraneuronal identity of these solitary cells.

On confocal images, the HCS-cells are round cells of approx. 5 μm in diameter located immediately underneath the pavement cell layer (the superficial layer) in the epidermis. Unlike keratinocytes, the HCS-cells did not remarkably display immunoreactivity with cytokeratin and desmosome antibodies. Interestingly, the HCS-cells did not display the characteristic features of hair cells and chemosensory cells, as shown by DASPEI staining and electron microscopy. Furthermore, ultrustructural analysis indicated that these cells lacked the classical hallmarks of tactile sensors called Merkel cells, such as dense-core granules, finger-like cytoplasmic processes, lobulated nucleus and an oval shape (Halata et al., 2003). However, the common feature with certain tactile sensors is the 5-HT-immunoreactivity, which was reported in Merkel-cells of touch domes and upper hair follicle of several mammalian species (García-Caballero et al., 1989; English et al., 1992). Merkel cells are touch sensitive cells with Piezo2-dependent transduction channels (Woo et al., 2015). They show immunoreactivity not only for serotonin, but also for a variety of neuromodulators, including vasoactive intestinal polypeptide, met-enkephalin, calcitonin gene-related peptide, somatostatin (Halata et al., 2003). However, there is no clear evidence to support a transmitter role for any of these substances (Halata et al., 2003). In zebrafish, 5-HT-immunoreactivity was shown in Merkel-like basal cells of the taste bud (Zachar and Jonz, 2012; Larson et al., 2015). On the other hand, Calretinin has not been reported in the basal cells of the taste buds, but in the chemosensory cells instead (Soulika et al., 2016; Ikenaga and Kiyohara, 2018). Thus, the exact function of the HCS-paraneurons remains to be identified in the future.

The biosynthesis of 5-HT requires the enzyme Tryptophan hydoroxylase (Tph), which catalyzes the rate-limiting step of the process (Lv and Liu, 2017). In mammals, *tph1* is typically expressed in peripheral tissues for serotonin synthesis, while *tph2* is expressed in the nervous system. Surprisingly, our *in-situ* hybridization detected the expression of *tph2* in the dispersed cells of the epidermis. In mammals, *tph2* transcripts have also been detected in certain peripheral cells, such as the retinal pigment epithelium and the aorta-gonad-mesonephrons (Żmijewski et al., 2009; Lv et al., 2017). Furthermore, rodent taste buds express both *tph1* and *tph2* (Dvoryanchikov et al., 2007). Overall, HCS-cells appear to express *tph2* for the local synthesis of serotonin.

Serotonin can act through receptor-dependent and -independent mechanisms, as a neurotransmitter, hormone, cytokine, biological modifier, growth factor, morphogen and anti-oxidant (Azmitia, 2001). In the mammalian skin, this monoamine plays diverse para- and endocrine roles by regulating cell proliferation, metabolism, resistance to stress, and immune reactions (Slominski et al., 2005). Based on this knowledge, it is possible that the 5-HT produced by HCS-cells acts not only as a neural messenger, but also as a local modulator in the epidermis. Further functional analyses are required to address these hypotheses.

### Distribution and regeneration of HCS-cells in the fin

The average density of HCS-cells was approx. 140 cells per mm^2^, with considerable variation even between sibling zebrafish raised in the same tanks. This variation might depend on individual swimming activities influenced by their hierarchal status or other behavioral cues. Irrespectively of average densities, the distribution pattern of the HCS-cells of both wild type and *alf* mutant fins revealed an increased density of these cells at ray bifurcation sites. During fin regeneration after amputation, the density was reproduced concomitantly with the new tissue. After completion of regeneration, the distribution pattern was reproduced, suggesting its functional importance. Two proliferation assays, based on PCNA expression and BrdU incorporation, suggested that mature fin HCS-paraneurons are not directly mitotic cells. This result is consistent with the individual distribution pattern of these cells, which do not form colonies. They likely arise by asymmetric cell division followed by a slow differentiation, a common mechanism for generation of neurosensory organs (Kapsimali, 2017). Furthermore, by blocking the enzymatic activity of Tph, we showed that serotonin synthesis is not necessary for fin regeneration and restoration of HCS-cells.

### Putative hydrodynamic sensing by paraneuronal cells

The spatial distribution of HCS-cells, characterized by higher densities inside the bifurcation interrays, is correlated to specific hydrodynamic conditions. A detailed analysis of the flow patterns in these regions, where the V-shaped obstacle causes a more turbulent boundary layer, supports the hypothesis that the HCS-cells might have a sensory function. In this scenario, the spatial distribution of HCS-cells is related to the capacity of encoding shear stress levels and turbulent fluctuations.

To this day, hydrodynamic sensing by aquatic animals has mainly been studied in the context of oscillatory signals in the fluid, traveling from a source distant from the fish, such as a prey or a predator. This type of hydrodynamic sensation is achieved by the canal neuromasts of the lateral line and the free neuromasts scattered across the body (McHenry et al., 2008; Van Trump and McHenry, 2008; Monroe et al., 2015). In contrast to the perception of distant signals, we postulated that local hydrodynamic changes in the vicinity of the caudal fin, such as shear stress amplitude and fluctuations due to turbulence in the boundary layer, could be detected by the HCS paraneuronal cells. Hence, this hydrodynamic analogue to tactile sensation would provide the fish with mechanical feedback from the surrounding water, allowing the fine-tuning of fin deformation for precise maneuvering. This mechanical feedback from the water, possibly combined with proprioception, would provide the animal with a rich set of information on which to base the coordination of its caudal fin relative to the fluid.

### Flow disturbances and turbulence induced in the bifurcation interray

The presence of a small obstacle on the surface of a propulsive appendage can induce the transition to turbulence inside a boundary layer (Anderson et al., 2001). Significant turbulence effects can be observed with 2D obstacles with a ratio of height to boundary layer thickness (*h*/*δ*) around 0.07-0.13 or higher (Arnal, 1984). To evaluate the extent to which the bifurcated rays of our model and of the real caudal fin could promote turbulence, we calculate the laminar boundary layer thickness over a flat plate (Schlichting and Gersten, 2017):

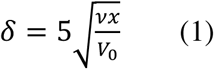

The distance *x* is measured from the edge of the plate (25 mm) or from the caudal fin peduncle (∼3.5 mm), and *V*_0_ is the free-stream velocity, equal to 55 mm/s in our hydrodynamic experiments, and 400 mm/s in the fastest case for a swimming zebrafish (Puri et al., 2017) (Fig. 9E). This results in an approximate boundary layer thickness of 3.4 mm for the plate model, and 0.47 mm for the caudal fin. The boundary layer thickness measured on the stationary non-bifurcated rays model shows good agreement with that theoretical value (Fig. 12A; top panel, triangles). The height of the obstacle for the plate model is the radius of the bifurcated rod, ∼0.88 mm. For the caudal fin, it is the height of the hemiray bulging out of the fin membrane, i.e. approximately 90 *μ*m (Murciano et al., 2018). The ratio between the obstacle height and the boundary layer thickness (*h*/*δ*) was 0.26 for the plate model, and 0.19 for the real caudal fin. This indicates that turbulent effects should be induced by the bifurcated rods placed across the path of the incoming flow.

An important notion to consider is the difference in thickness between the primary interray and the bifurcation interray. According to the previous measurements (Murciano et al., 2018), the paraneuronal cells located inside the bifurcation interray are placed further inside the flow than those located in the adjacent primary interrays, with a difference in distance of the order of 100 μm. This abrupt change in surface level places the paraneuronal cells of the bifurcation interray in a different portion of the external flow, possibly granting them access to clearer, easier to encode, shear stress signals, related to stronger turbulence.

Taken together, the HCS paraneuronal cells are distributed in a way that correlated with fluid dynamics. The existence of these cells highlights the diversity of cells present in the zebrafish skin and emphasizes how the aquatic environment impacts the way in which neurosecretory modalities can be distributed on the body.

## Materials and methods

### Fish strains and fin amputations

For this study, the following strains were used: wild-type AB strain (Oregon), *alf^dty86^* (*another longfin^dty86^*) (Perathoner et al., 2014). Adult fish were used at ages 10-24 months, larvae were used at 14 days-post fertilization and juvenile fish at 21-60 days post-fertilization. For fin amputation, fish were anesthetized in 0.6 mM tricaine (MS-222 ethyl-m-aminobenzoate, Sigma-Aldrich), and fins were amputated with a razor blade proximal to the first bone bifurcation point. Animals were kept at 27°C for various durations before fin collection. Live images (bright field) of fins were taken with a Leica AF M205 FA stereomicroscope. Animal experimentation was approved by the Cantonal Veterinary office of Fribourg, Switzerland.

### Drug treatments

Drug treatments were performed in 100 mL coplin jars (up to 3 fish per coplin jar). The following drugs were used: para-chlorophenylalanin (pcpa, 5 mM; Sigma-Aldrich), BrdU (163 µM; Sigma-Aldrich). Control fish were kept in 100 mL coplin jars.

### Whole-mount *in situ* hybridization

For the detection of transcripts in the fin epidermis, Zinc-fixation followed by whole mount *in situ* hybridization were performed with a protocol adapted from (Stylianopoulou et al., 2014).

Uninjured adult fins were collected in TS (0.1 M Tric-HCl pH7, 0.05 M NaCl), rinsed in TS and fixed in Z7 zinc fixative (0.5% Zinc choride, 0.5% Zinc trifluoroacetate, 0.05% calcium acetate, 0.1 M Tris-HCl pH7, filtered) for 15 minutes. They were then rinsed in TS and post-fixed in 4% Paraformaldehyde for 10 minutes. Fins were washed twice for 5 minutes in PBS and permeabilized, first in 1.3 % v/v triethanolamine, 0.03 N HCl, 0.25% acetic anhydride for 10 minutes, then in PBS + 1% Triton-X for 30 minutes. Fins were washed again in PBS twice and transferred to spot plates.

In spot plates, fins were incubated in hybridization solution (50% formamide, 5x SSC, 10% Dextran, 0.1 mg/mL tRNA) for 10 min at RT and for 1 h at 60°C. The fins were then incubated overnight at 60°C with dig-labeled RNA antisense probes diluted in hybridization solution (Spot plate wells were covered with plastic coverslips to prevent evaporation). On the next day, the fins were washed in decreasing dilution series of hybridization solution in 5xSSC for 10 minutes each, 5x SSC for 10 min and 0.2x SSC for 30 minutes twice.

For antibody labeling, the fins were incubated in increasing dilution series of maleic acid buffer (100 mM maleic acid, 150 mM NaCl, pH 7.5) in 0.2x SSC for 10 minutes each to reach 100% maleic acid buffer. Blocking was performed for 1 h in blocking solution (Roche) diluted in maleic acid buffer. The fins were then incubated with anti-dig antibody in blocking solution (1:4000, Roche) for 2 h. After incubation, fins were washed for 30 minutes in maleic acid buffer.

For colorimetric reaction, the fins were washed twice 5 min in AP buffer (1M Tris-HCl pH 9.5, 1 M MgCl_2_, 5 M NaCl, 20% Tween-20). The colorimetric reaction was then induced by incubation in AP buffer + 0.2 mg/mL NBT + 0.3 mg/mL BCIP. The reaction was performed at 37°C. The fins were monitored every 10 minutes and the reaction was stopped by transferring fins to PBS for 15 minutes followed by 70% ethanol for 1h. The fins were mounted in 80% glycerol mounting medium and imaged immediately.

The following Primers were used for the generation of dig-labeled antisense probes: *tph1a (NM_178306.3): Fw 5’-aagcgagatggagaatgtgc-3’; Rev: 5’-tgcatctccaagatgtccag-3’ tph1b (NM_001001843.2): Fw 5’-gggctggtcttctctcttcc-3’; Rev: 5’-cagctcatggcaagaaacag-3’ tph2 (NM_001310068.1): Fw 5’-tctcagagctggatcagtgc-3’; Rev: 5’-tcacagacggtggttagtcg-3’*

### Electron microscopy

For Transmission electron microscopy (TEM), samples from four fish have been analyzed. Caudal fins of wild type zebrafish at the age of 14 months were processed according to standard protocol. Briefly, the fins were fixed in Karnovsky’s solution, washed in sodium cacodylate buffer, dehydrated in a graded series of ethanol, and embedded in EPON resin (Sigma-Aldrich). Blocks were cut at the thickness of 60 nm with an ultra-microtome (Leica, Germany), equipped with a diamond knife (Diatome, Switzerland). Sections were mounted on copper specimen grids (Plano, Germany) and stained with uranyl acetate and lead citrate for 40 min. The specimens were examined using a Philips TEM CM12 electron microscope. Two to three paraneurons were imaged per sample.

For scanning electron microscopy (SEM), three caudal fins were processed according to standard procedure. Briefly, the specimens were fixed in Karnovsky solution, washed in sodium cacodylate buffer, dehydrated in increasing concentrations of ethanol in water. Dehydrated samples were dried by the critical point drying method, using liquid carbon dioxide in a pressure chamber and mounted on aluminum stubs. The specimens were sputter-coated with gold and viewed under a Philips XL 30 FEG scanning electron microscope.

### Immunofluorescence staining (sections and whole-mount)

Immunofluorescence analyses of fin sections were performed as previously described (König et al., 2018). Briefly, fins were harvested, fixed in 2% paraformaldehyde (PFA) overnight at 4°C and embedded in tissue freezing medium. Sections were cut to 16 µm thickness. Two step immunofluorescence staining was performed.

For whole-mount analysis, adult fins or full larvae were harvested and fixed in 2% PFA overnight at 4°C (Thorimbert et al., 2015). They were washed in phosphate-buffered saline (PBS) after fixation, transferred to blocking solution (5% goat serum in PBS+0.3% Triton) for one hour. They were then incubated in a primary antibody diluted in blocking solution overnight at 4°C. On the next day, fins were washed and incubated with a secondary antibody diluted in blocking solution for 2 h at room temperature. All the steps were done on a rocking platform. The fins were washed again and mounted. For BrdU visualization, the fins were incubated for 25 minutes in a solution of 2 N HCl before immunofluorescence staining.

The following primary antibodies were used: Rabbit anti-serotonin (1:2000; Sigma-Aldrich; S5545; immunogen: serotonin creatinine sulfate complex conjugated to BSA). This antibody gives a very specific and strong staining in immunofluoresce assays. Rat monoclonal antibody (clone YC5/45) anti-serotonin (1:50; Abcam; ab6336; immunogen: serotonin conjugated to BSA) and Mouse monoclonal antibody against serotonin (1:10; Novusbio; NB120-16007). These antibodies are weaker and give more background as compared to the rabbit anti-serotonin antibody; they did not work for whole mount immunolabeling. Mouse monoclonal (clone Zn-12) antibody anti-L2/HNK-1 tetrasaccharide (1:100; ZFIN); Rabbit anti-Calretinin (1:1000; 7699/4; Swant. This antibody was validated in zebrafish by Western Blot and immunofluorescence (Castro et al., 2006)); Mouse anti-SV2 (1:50; Synaptic vesicle glycoprotein 2; DSHB); Mouse anti-keratin (1:500; USBiological; K01199-90; pan antibody cocktail containing AE1/AE3 clones; recognizes acidic (Type I or LMW) and basic (Type II or HMW) cytokeratins); Mouse anti-desmoplakin 1/2 (1:50; Progen); Rabbit anti-tp63 (1:500; Genetex); Rabbit anti-PCNA (1:200; Genetex; GTX124496); Rat anti-BrdU (1:250; Abcam; ab6326). Fluorescent dye-coupled secondary antibodies (Jackson Immunological) were used at 1:500. DAPI (Sigma-Aldrich) was used to label nuclei. Immunofluorescence was imaged using a Leica SP5 confocal microscope.

### Image processing and analysis

Images from confocal microscopy were processed using Image J (NIH) and Photoshop (Adobe).

Overall density of HCS-cells was calculated in Image J as follows: The 2^nd^-7^th^ rays from the lateral edge were traced in Image J and the area was measured (in mm^2^). In order to count the 5-HT-positive cells (or SV2-, Calretinin- or Tp63-positive cells) a threshold was applied on the 5-HT channel (Analyze > Threshold); this eliminates background signal and autofluorescence of pigments, and turns the image to binary. The number of cells was then measured using the ‘Analyze particles’ tool (Setting: 10-100 µm^2^, 0-1 circularity). The density of cells in the epidermis was calculated by dividing the number of cells by the area and by two (in order to show the density for each epidermal surface). When quantifying density in regenerating fins, only the area beyond the amputation plane was taken into account.

Density of HCS-cells in primary interray (PI) versus bifurcation interray (BI) was calculated as follows: the primary bifurcations of rays 3, 4 and 5 were considered. Areas between bifurcations (BI) were selected in a range of 5 bone segments distal from the bifurcation point. Corresponding primary interray (PI) areas were selected at the same proximo-distal level adjacent to the bifurcation (Fig. 6C). For each of the 6 selected zones (3 PI and 3 BI), areas were measured and numbers of serotonin-positive cells in the area quantified as described above. For each fin, the sum of all cells in the three BI was divided by the sum of the three BI areas, respectively all the cells in adjacent PI divided by the area of the selected PI zones.

Projections of 5-HT-positive cells were performed as follows: The positions of the 5-HT-positive cells were obtained by repeating the above procedure in Image J on whole images (threshold on serotonin channel followed by particle analysis), but the analyse particles tool was used with the ‘record start’ option selected. The output is a list of cells with their sizes and their exact positions (X-Y coordinates). This was transferred to Microsoft Excel where a scatter plot was created using the coordinates (white dots on transparent background). The scatterplot was then overlaid over the image of the fin in Photoshop (matching the dots of the scatterplot to the 5-HT-positive cells in the image).

Percentage of BrdU positive cells was counted on optical confocal sections through the epidermis of a whole mount image. The manual cell counter tool was used in Image J. Each 5-HT-positive cell was assigned to be either BrdU-positive or –negative. The proportion of BrdU-positive HCS-cells was then calculated. Percentage of PCNA positive cells was analyzed similarly, assigning all SV2-positive HCS-cells as either PCNA-positive or –negative. Adjacent SV2-negative epidermal cells were used for counting the proportion of SV2-negative/PCNA-positive cells.

Graphs were plotted in Graphpad prism. Statistical analyses were performed using student’s T tests. Normal distribution of the data sets was confirmed using the Shapiro-Wilk normality test. When comparing overall density between different groups of fish (transgenes vs wild-type, treated vs control, different ages), unpaired T-tests were done. When comparing density in bifurcation interray versus primary interrays, paired T-tests were used. Unless otherwise specified Means are indicated ± standard deviation.

### DASPEI labeling

2-[4-(Dimethylamino)styryl]-1-ethylpyridinium iodide (DASPEI) labeling in live fish was performed by diluting 1mM DASPEI (Sigma-Aldrich) in system water. Fish were incubated in the solution for 1 minute, quickly rinsed in system water and anesthetized for imaging. Images were taken with a Leica AF M205 FA stereomicroscope with the GFP2 filter. DASPEI cells were counted manually in ImageJ using the cell counter tool, because the tissue was too autofluorescent for quantification through the analyze particle option.

### Analysis of fluid dynamics on bifurcation model

In order to study and compare the specific hydrodynamic profiles inside the bifurcation and the primary interrays, a simplified model consisting of a rigid plate supporting half-cylindrical rods was designed, with a control version simply made of three parallel rods and a second version including a bifurcation in the central ray. The geometry and dimensions of the two plate models are illustrated in Fig. 9C (all dimensions in mm). The plate models were consecutively mounted inside a flow chamber and actuated with a servomotor fixed below the tunnel to mimic the oscillatory motion of the zebrafish caudal fin, as seen in Fig. 9, where a front and a top view of the plate model mounted inside the water tunnel are shown (Ba and Bb). Additionally to the flapping motion, an upstream constant flow speed was maintained by an external pump. The model dimensions and kinetic aspects of the flow were scaled appropriately to obtain values of the Reynolds and Strouhal numbers in the same range as those corresponding to the zebrafish caudal fin during typical cruising (Fig. 9E). This proper dimension scaling ensures that the model operates in a similar hydrodynamic regime as the imitated system. The Reynolds number encompasses the ratio of inertial over viscous forces and determines the transition from a laminar to a turbulent flow, whereas the Strouhal number describes the propulsion dependence on the tail oscillation:

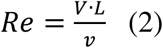

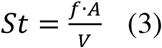

The zebrafish exhibits a large variety of swimming behaviors, and therefore covers up a wide range of swimming speed, tail beat frequency and amplitude. The values displayed in (Fig. 9E) are derived from (Palstra et al., 2010; Mwaffo et al., 2017), and are presented as an approximation of the caudal fin dynamics during the cruising propulsion mode. An estimate of the caudal fin length is used to calculate the Reynolds number so as to remain in the framework of the plate model, which explains a lower range than what is usually obtained when the whole fish size is used as the characteristic length in equation (2). The model oscillation parameters (period ≅ 2 s, *θ*_0_ ≅ 11°) yield a Strouhal number of about 0.15, which resides in a suboptimal range in terms of animal propulsion efficiency (Fig. 9E). Nevertheless, it is probable that the caudal fin dynamics are different during chemotaxis-type behaviors as compared to efficient swimming.

The hydrodynamic experiment is achieved using a three-dimensional particle tracking velocimetry set up from TSI, Inc. (V3V-9800-TS) as described in http://www.tsi.com/V3V-9800-System/. This system allows the measurement of instantaneous velocity flow fields surrounding an object in motion, based on the seeding of tracer particles (∼50 μm polyamide) in the fluid volume, which we illuminate and image using a laser synchronized with three high-speed cameras. The velocity vectors of the advected tracers are subsequently computed from the image sequences using appropriate particle tracking algorithms. The basic principle behind particle imaging velocimetry (PIV) techniques for flow analysis is the detection of tracer particles and the reconstruction of their positions inside the measurement volume. By computing the distance traveled by each tracer particle between subsequent time frames, and by setting a fixed time duration between the image captures, the particles instantaneous velocities (Δ*x*/Δ*t*) can be calculated, thus allowing the reconstruction of the whole fluid velocity field. For an exhaustive description of the PIV method, the reader is referred to (Raffel et al., 2018). The experimental set up and its various components (flow chamber and PIV interrogation volume, dual-head pulsed laser, camera triplet, water reservoirs and pumping mechanism) are illustrated and described in Fig. 9.

Each of the three cameras captures an image of the interrogation volume from a different perspective, allowing the particle positions to be reconstructed including the third dimension (depth). In Fig. 9, an example is provided of a triplet of raw images, each recorded by a distinct camera (left, right and top cameras mounted on a triangular plate). The plate is visible in those images, owing to equally spaced white dots which were painted directly on the half-cylindrical rods. Using a specific set of parameters, the images can be preprocessed in order to detect these dots only, and to locate precisely the position of the oscillating plate inside the flow chamber at each analyzed motion instant. Nevertheless, at all instants of its oscillation cycle, the plate remains inside the PIV interrogation volume. The plate was fabricated using a matte black rigid plastic (polyoxymethylene) in order to reduce reflections from the laser beam and to allow the detection of tracer particles almost up to the surface itself, which is necessary to resolve the boundary layer properly. Thus, all particles recorded by the cameras are located in front of the plate. The number of particles velocity vectors reconstructed in the region located in front of the plate (about half the interrogation volume) is ∼3200 (after applying the median and range filters described below). In total, ∼30 *mL* of particles were seeded in ∼160 *mL* of water, which yields an approximate number of 7200 particles inside the 50×50×20 *mm*^2^ PIV interrogation volume, considering that the spherical particles powder has a filling fraction of ∼40%.

Uncertainties on the detected particle positions in each of the triplet images, namely the X and Y positions, depend on a number of experimental factors such as the particle size, the cameras magnification, the quality of the calibration, etc. Combining those within the error propagation method results in a positional uncertainty of *σ_x_* = *σ_y_* ≅ 3.6 μm for the in-plane components, which in turns, according to the geometrical arrangement of the cameras lenses, yields an uncertainty of *σ_z_*≅ 32 μm for the out-of-plane component. The error propagation then leads to equation (4) for the velocity components uncertainties: *σ_v_x__* = *σ_v_y__*≅ 0.6 mm/s and *σ_v_z__* ≅ 5 mm/s (with Δt = 0.009 s).

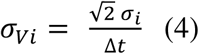

Appropriate filters (median filter / *ad hoc* range for velocity vectors) were used to reduce the data noise (outlier vectors). Additionally, temporal averaging was performed for each position of the plate (three to five timeframes for each position). Consequently, the values calculated above are reduced by at least a factor of two. In addition, the velocities in Fig. 9E present averages of the flow over a relatively large area, where a large number of tracer particles are averaged over. This leads to a further reduction of the uncertainty on the average by 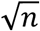, where n is the number of particles in the averaging region. Therefore, we estimate an uncertainty of 0.2 mm/s for the average parallel flow speeds and of 1.0 mm/s for the average perpendicular flow speeds.

The flow measurement was conducted over a complete period of oscillation of the plate model, and for purpose of conciseness, the fluid velocity profiles were extracted at four key positions, equally distributed through the full cycle, and presented in order in Fig. 11A) traveling (with maximum velocity magnitude) through the 0° angle towards the front, B) instantly immobile at the maximum angle, C) traveling again through the 0° angle but towards the back, and D) minimum angular position which also corresponds to the back of the flow chamber. For each of these instants, we focused our attention on vertical planes extracted from the 3D velocity fields which are parallel and directly adjacent to the plate model. In other words, the 2D slices from the 3D velocity fields presented in Fig. 10 and analyzed in Fig 11 correspond to vertical planes directly adjacent and parallel to the plate model. Due to the varying angle of the plate over the oscillation period, the velocity vectors are decomposed based on a coordinate system attached to the plate (rather than a fixed cartesian coordinate system). In other words, the fluid velocity is decomposed into the following three orthogonal components: one parallel to the plate and noted *V*_║_ (positive values indicate that the fluid is moving in the upstream direction), one perpendicular to the plate and noted *V*_⊥_, (positive values denote fluid motion away from the plate and vice-versa) and a vertical component noted *V*_+_ (positive values refer to an upward fluid motion). Moreover, all velocity components are evaluated in the plate’s referential, which means that they are expressed relatively to the plate’s velocity. The relevant axes are illustrated in Fig. 9Ba and Bb.

The decomposition of fluid velocity vectors into the appropriate components at specific instants of motion allows the visualization of interesting flow signatures in the inter-bifurcation region. The exact quantification of these flow features is done by averaging each velocity component over two distinct area types, namely inside the V-shape bifurcation (bifurcation interray, noted BI) and between the adjacent neighboring rays (primary interray, noted PI) (Fig. 9C).

The boundary layer that forms over the surface of an immerged object originates from the no-slip condition, which states that the fluid velocity relative to the solid is null at the interface. Consequently, the fluid velocity increases rapidly from zero to the free-stream value in order to satisfy this boundary condition. The streamwise velocity far away from the object is defined in our experiment as *V*_║_*_ave_*, the average value of *V*_║_ over the whole measurement volume, at each instant of motion. The thin region over which this velocity change occurs is called the boundary layer. Its thickness is defined as the distance, measured from the solid surface, at which the velocity has reached 99% of the free-stream value:

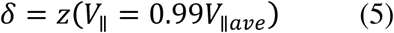

The boundary layer thickness was evaluated from the PIV velocity fields, by extracting horizontal planes from the measurement volume at locations of interest. The velocity vectors were averaged for three specific regions: (i) between the parallel rods of the non-bifurcated rays plate model, (ii) in the primary interray of the bifurcated ray model, and (iii) in the bifurcation interray of that same model. For the first case, 18 equidistant *x* − *z* planes were used for the average, 11 for the PI region, and 4 for the BI zone. This process was repeated for three different motion instants: one where the plate was held stationary, and the 0° angle position of the plate during the pitching motion, both for the forward and the backward portion of the oscillation cycle. This resulted in a 2D top view of the flow for every region and time frame. In each case, the streamwise component of the velocity *V*_║_ was normalized by the average value *V*_║*ave*_ and illustrated with a color map (Fig. 12B). Subsequently, the boundary layer thickness was evaluated as the *z* distance measured from the plate at which the velocity reached 99% of *V*_║*ave*_ (Fig. 12A).

## Acknowledgement

We are grateful to V. Zimmermann and J. Wagner for technical help and animal care support, T. Aegerter-Wilmsen for interdisciplinary discussions, and C. Pfefferli for critical reading of the manuscript.

## Author contribution

DK designed and carried out experiments with fish, analyzed the biological data and contributed to writing the manuscript. PD and CA designed and carried out kinematic experiments, and contributed to writing the manuscript. AS and VD designed and carried out electron microscopy experiments. AJ designed the experiments and wrote the manuscript.

## Funding

This work was supported by the Swiss National Science Foundation, project grant: 310030_179213 and Sinergia grant CRSII3_147675.

## Competing interest

The authors declare no financial or non-financial competing interests.

**Supplementary Figure. Images of Zooms from main figures without white cell-projection dots.** The corresponding main figures are indicated above the image. Tissue Autofluorescence in pink. Serotonin-positive cells in green.

## References

Aiello, B.R., Westneat, M.W., and Hale, M.E. (2017). Mechanosensation is evolutionarily tuned to locomotor mechanics. Proceedings of the National Academy of Sciences 114(17), 4459–4464. doi: 10.1073/pnas.1616839114.

Airhart, M.J., Lee, D.H., Wilson, T.D., Miller, B.E., Miller, M.N., Skalko, R.G., et al. (2012). Adverse effects of serotonin depletion in developing zebrafish. Neurotoxicology and Teratology 34(1), 152–160. doi: 10.1016/j.ntt.2011.08.008.

Akimenko, M.A., Mari-Beffa, M., Becerra, J., and Geraudie, J. (2003). Old questions, new tools, and some answers to the mystery of fin regeneration. Dev Dyn 226(2), 190–201. doi: 10.1002/dvdy.10248.

Anderson, E.J., McGillis, W.R., and Grosenbaugh, M.A. (2001). The boundary layer of swimming fish. Journal of Experimental Biology 204(1), 81–102.

Arnal, D. (1984). Description and Prediction of Transition in Two-Dimensional, Incompressible Flow. OFFICE NATIONAL D’ETUDES ET DE RECHERCHES AEROSPATIALES TOULOUSE: Defense Technical Information Center.

Atema, J., Fay, R.R., Popper, A.N., and Tavolga, W.N. (1988). Sensory Biology of Aquatic Animals. New York: Springer.

Azmitia, E.C. (2001). Modern views on an ancient chemical: serotonin effects on cell proliferation, maturation, and apoptosis. Brain Research Bulletin 56(5), 413–424. doi: 10.1016/s0361-9230(01)00614-1.

Berger, M., Gray, J.A., and Roth, B.L. (2009). The Expanded Biology of Serotonin. Annual Review of Medicine 60(1), 355–366. doi: 10.1146/annurev.med.60.042307.110802.

Bleckmann, H., Mogdans, J., and Coombs, S.L. (2014). Flow Sensing in Air and Water: Behavioral, Neural and Engineering Principles of Operation. Berlin Heidelberg: Springer Science & Business Media.

Buckley, K., and Kelly, R.B. (1985). Identification of a transmembrane glycoprotein specific for secretory vesicles of neural and endocrine cells. The Journal of Cell Biology 100(4), 1284–1294. doi: 10.1083/jcb.100.4.1284.

Castro, A., Becerra, M., Manso, M.J., and Anadón, R. (2006). Calretinin immunoreactivity in the brain of the zebrafish, Danio rerio: Distribution and comparison with some neuropeptides and neurotransmitter-synthesizing enzymes. I. Olfactory organ and forebrain. The Journal of Comparative Neurology 494(3), 435–459. doi: 10.1002/cne.20782.

Coccimiglio, M.L., and Jonz, M.G. (2012). Serotonergic neuroepithelial cells of the skin in developing zebrafish: morphology, innervation and oxygen-sensitive properties. Journal of Experimental Biology 215(22), 3881–3894. doi: 10.1242/jeb.074575.

Collin, S.P., Bullock, T., Atema, J., Marshall, N.J., Fay, R.R., Popper, A.N., et al. (2008). Sensory Processing in Aquatic Environments. Springer New York.

Coombs, S., Bleckmann, H., Fay, R., and Popper, A. (2013). The Lateral Line System. Springer Science & Business Media.

Delva, E., Tucker, D.K., and Kowalczyk, A.P. (2009). The Desmosome. Cold Spring Harbor Perspectives in Biology 1(2), a002543–a002543. doi: 10.1101/cshperspect.a002543.

Doughty, M.J., Bergmanson, J.P.G., and Blocker, Y. (1997). Shrinkage and distortion of the rabbit corneal endothelial cell mosaic caused by a high osmolality glutaraldehyde-formaldehyde fixative compared to glutaraldehyde. Tissue and Cell 29(5), 533–547. doi: 10.1016/s0040-8166(97)80054-7.

Dufourcq, P., Roussigné, M., Blader, P., Rosa, F., Peyrieras, N., and Vriz, S. (2006). Mechano-sensory organ regeneration in adults: The zebrafish lateral line as a model. Molecular and Cellular Neuroscience 33(2), 180–187. doi: 10.1016/j.mcn.2006.07.005.

Dvoryanchikov, G., Tomchik, S.M., and Chaudhari, N. (2007). Biogenic amine synthesis and uptake in rodent taste buds. The Journal of Comparative Neurology 505(3), 302–313. doi: 10.1002/cne.21494.

English, K.B., Wang, Z.-Z., Stayner, N., Stensaas, L.J., Martin, H., and Tuckett, R.P. (1992). Serotonin-like immunoreactivity in Merkel cells and their afferent neurons in touch domes from the hairy skin of rats. The Anatomical Record 232(1), 112–120. doi: doi:10.1002/ar.1092320112.

Finger, T.E. (2009). “Evolution of taste systems.,” in Evolutionary neuroscience, eds. J.H. Kaas, G.F. Striedter, J. Rubenstein, T.H. Bulluck, L. Krubitzer & T. Preuss. 1st ed (Oxford, UK: Academic Press), 460–477.

Flammang, B.E., and Lauder, G.V. (2013). Pectoral fins aid in navigation of a complex environment by bluegill sunfish under sensory deprivation conditions. The Journal of Experimental Biology 216(16), 3084–3089. doi: 10.1242/jeb.080077.

Foskett, J.K., Logsdon, C.D., Turner, T., Machen, T.E., and Bern, H.A. (1981). Differentiation of the Chloride Extrusion Mechanism During Seawater Adaptation of a Teleost Fish, The Cichlid *Sarotherodon Mossambicus*. Journal of Experimental Biology 93(1), 209.

Froehlicher, M., Liedtke, A., Groh, K.J., Neuhauss, S.C.F., Segner, H., and Eggen, R.I.L. (2009). Zebrafish (Danio rerio) neuromast: Promising biological endpoint linking developmental and toxicological studies. Aquatic Toxicology 95(4), 307–319. doi: 10.1016/j.aquatox.2009.04.007.

Fujita, T. (1989). Present status of paraneuron concept. Archives of Histology and Cytology 52(Suppl), 1–8. doi: 10.1679/aohc.52.Suppl_1.

Fujita, T., Kanno, T., and Kobayashi, S. (1988). The paraneuron. Tokyo: Springer Japan.

García-Caballero, T., Gallego, R., Rosón, E., Basanta, D., Morel, G., and Beiras, A. (1989). Localization of serotonin-like immunoreactivity in the merkel cells of pig snout skin. The Anatomical Record 225(4), 267–271. doi: doi:10.1002/ar.1092250402.

Ghysen, A., and Dambly-Chaudiere, C. (2007). The lateral line microcosmos. Genes & Development 21(17), 2118–2130. doi: 10.1101/gad.1568407.

Goldsmith, M.I., Iovine, M.K., O’Reilly-Pol, T., and Johnson, S.L. (2006). A developmental transition in growth control during zebrafish caudal fin development. Dev Biol 296(2), 450–457. doi: 10.1016/j.ydbio.2006.06.010.

Halata, Z., Grim, M., and Bauman, K.I. (2003). Friedrich Sigmund Merkel and his “Merkel cell”, morphology, development, and physiology: Review and new results. The Anatomical Record Part A: Discoveries in Molecular, Cellular, and Evolutionary Biology 271A(1), 225–239. doi: doi:10.1002/ar.a.10029.

Hardy, A.R., Steinworth, B.M., and Hale, M.E. (2016). Touch sensation by pectoral fins of the catfishPimelodus pictus. Proceedings of the Royal Society B: Biological Sciences 283(1824), 20152652. doi: 10.1098/rspb.2015.2652.

Iger, Y., and Wendelaar Bonga, S.E. (1994). Cellular responses of the skin of carp (Cyprinus carpio) exposed to acidified water. Cell and Tissue Research 275(3), 481–492. doi: 10.1007/bf00318817.

Ikenaga, T., and Kiyohara, S. (2018). “Chemosensory Systems in the Sea Catfish, Plotosus japonicus,” in Zebrafish, Medaka and Other Small Fishes, eds. H. Hirata & A. Iida. (Singapore: Springer), 295–315.

Jaźwińska, A., Badakov, R., and Keating, M.T. (2007). Activin-betaA signaling is required for zebrafish fin regeneration. Curr Biol 17(16), 1390–1395.

Jequier, E., Lovenberg, W., and Sjoerdsma, A. (1967). Tryptophan hydroxylase inhibition: the mechanism by which p-chlorophenylalanine depletes rat brain serotonin. Mol Pharmacol 3(3), 274–278.

Jørgensen, J.M. (1992). The Electrosensory Cells of the Ampullary Organ of the Transparent Catfish (Kryptopterus bicirrhus). Acta Zoologica 73(2), 79–83. doi: doi:10.1111/j.1463-6395.1992.tb00951.x.

Kapsimali, M. (2017). Epithelial cell behaviours during neurosensory organ formation. Development 144(11), 1926–1936. doi: 10.1242/dev.148122.

Kasumyan, A.O. (2011). Tactile reception and behavior of fish. Journal of Ichthyology 51(11), 1035–1103. doi: 10.1134/s003294521111004x.

Kniss, J.S., Jiang, L., and Piotrowski, T. (2016). Insights into sensory hair cell regeneration from the zebrafish lateral line. Current Opinion in Genetics & Development 40, 32–40. doi: 10.1016/j.gde.2016.05.012.

König, D., and Jaźwińska, A. (2019). Zebrafish fin regeneration involves transient serotonin synthesis. Wound Repair and Regeneration. doi: 10.1112/wrr.12719.

König, D., Page, L., Chassot, B., and Jaźwińska, A. (2018). Dynamics of actinotrichia regeneration in the adult zebrafish fin. Developmental Biology 433(2), 416–432. doi: 10.1016/j.ydbio.2017.07.024.

Kotrschal, K. (1996). Solitary chemosensory cells: why do primary aquatic vertebrates need another taste system? Trends Ecol Evol 11(3), 110–114. doi: 10.1016/0169-5347(96)81088-3.

Kotrschal, K., Krautgartner, W.-D., and Hansen, A. (1997). Ontogeny of the Solitary Chemosensory Cells in the Zebrafish, Danio rerio. Chemical Senses 22(2), 111–118. doi: 10.1093/chemse/22.2.111.

Lane, B., and Whitear, M. (1977). On the occurrence of merkel cells in the epidermis of teleost fishes. Cell and Tissue Research 182(2), 235–246. doi: 10.1007/bf00220592.

Larson, E.D., Vandenbeuch, A., Voigt, A., Meyerhof, W., Kinnamon, S.C., and Finger, T.E. (2015). The Role of 5-HT3 Receptors in Signaling from Taste Buds to Nerves. Journal of Neuroscience 35(48), 15984–15995. doi: 10.1523/jneurosci.1868-15.2015.

Lv, J., and Liu, F. (2017). The Role of Serotonin beyond the Central Nervous System during Embryogenesis. Frontiers in Cellular Neuroscience 11. doi: 10.3389/fncel.2017.00074.

Lv, J., Wang, L., Gao, Y., Ding, Y.-Q., and Liu, F. (2017). 5-hydroxytryptamine synthesized in the aorta-gonad-mesonephros regulates hematopoietic stem and progenitor cell survival. The Journal of Experimental Medicine 214(2), 529–545. doi: 10.1084/jem.20150906.

McHenry, M.J., Strother, J.A., and van Netten, S.M. (2008). Mechanical filtering by the boundary layer and fluid–structure interaction in the superficial neuromast of the fish lateral line system. Journal of Comparative Physiology A 194(9), 795. doi: 10.1007/s00359-008-0350-2.

Metcalfe, W.K., Myers, P.Z., Trevarrow, B., Bass, M.B., and Kimmel, C.B. (1990). Primary neurons that express the L2/HNK-1 carbohydrate during early development in the zebrafish. Development 110(2), 491–504.

Moghadam, P., and Jackson, M. (2013). The Functional Significance of Synaptotagmin Diversity in Neuroendocrine Secretion. Frontiers in Endocrinology 4(124). doi: 10.3389/fendo.2013.00124.

Monroe, J.D., Rajadinakaran, G., and Smith, M.E. (2015). Sensory hair cell death and regeneration in fishes. Frontiers in Cellular Neuroscience 9(131). doi: 10.3389/fncel.2015.00131.

Montesinos, E., Esteve, I., and Guerrero, R. (1983). Comparison between direct methods for determination of microbial cell volume: electron microscopy and electronic particle sizing. Applied and Environmental Microbiology 45(5), 1651–1658.

Murciano, C., Cazorla-Vázquez, S., Gutiérrez, J., Hijano, J.A., Ruiz-Sánchez, J., Mesa-Almagro, L., et al. (2018). Widening control of fin inter-rays in zebrafish and inferences about actinopterygian fins. Journal of Anatomy 232(5), 783–805. doi: 10.1111/joa.12785.

Mwaffo, V., Zhang, P., Romero Cruz, S., and Porfiri, M. (2017). Zebrafish swimming in the flow: a particle image velocimetry study. PeerJ 5, e4041. doi: 10.7717/peerj.4041.

Njagi, J., Ball, M., Best, M., Wallace, K.N., and Andreescu, S. (2010). Electrochemical Quantification of Serotonin in the Live Embryonic Zebrafish Intestine. Analytical Chemistry 82(5), 1822–1830. doi: 10.1021/ac902465v.

Palstra, A.P., Tudorache, C., Rovira, M., Brittijn, S.A., Burgerhout, E., van den Thillart, G.E.E.J.M., et al. (2010). Establishing Zebrafish as a Novel Exercise Model: Swimming Economy, Swimming-Enhanced Growth and Muscle Growth Marker Gene Expression. PLoS ONE 5(12). doi: 10.1371/journal.pone.0014483.

Pangršič, T., Gabrielaitis, M., Michanski, S., Schwaller, B., Wolf, F., Strenzke, N., et al. (2015). EF-hand protein Ca^2+^ buffers regulate Ca^2+^ influx and exocytosis in sensory hair cells. Proceedings of the National Academy of Sciences 112(9), E1028–E1037. doi: 10.1073/pnas.1416424112.

Parichy, D.M., Elizondo, M.R., Mills, M.G., Gordon, T.N., and Engeszer, R.E. (2009). Normal table of postembryonic zebrafish development: Staging by externally visible anatomy of the living fish. Developmental Dynamics 238(12), 2975–3015. doi: 10.1002/dvdy.22113.

Perathoner, S., Daane, J.M., Henrion, U., Seebohm, G., Higdon, C.W., Johnson, S.L., et al. (2014). Bioelectric signaling regulates size in zebrafish fins. PLoS Genet 10(1), e1004080. doi: 10.1371/journal.pgen.1004080.

Pfefferli, C., and Jaźwińska, A. (2015). The art of fin regeneration in zebrafish. Regeneration 2(2), 72–83. doi: 10.1002/reg2.33.

Puri, S., Aegerter-Wilmsen, T., Jaźwińska, A., and Aegerter, C.M. (2017). *In-vivo* quantification of mechanical properties of caudal fins in adult zebrafish. The Journal of Experimental Biology, jeb.171777. doi: 10.1242/jeb.171777.

Raffel, M., Willert, C.E., F., S., Kähler, C., Wereley, S.T., and Kompenhans, J. (2018). Particle image velocimetry. A practical Guide.: Springer International Publishing AG.

Schlichting, H., and Gersten, K. (2017). Boundary-Layer Theory. New York: Spinger-Verlag.

Schwaller, B. (2014). Calretinin: from a “simple” Ca2+ buffer to a multifunctional protein implicated in many biological processes. Frontiers in Neuroanatomy 8. doi: 10.3389/fnana.2014.00003.

Slominski, A., Wortsman, J., and Tobin, D.J. (2005). The cutaneous serotoninergic/melatoninergic system: securing a place under the sun. The FASEB Journal 19(2), 176–194. doi: 10.1096/fj.04-2079rev.

Slominski, A.T., Zmijewski, M.A., Skobowiat, C., Zbytek, B., Slominski, R.M., and Steketee, J.D. (2012). Sensing the environment: regulation of local and global homeostasis by the skin’s neuroendocrine system. Advances in anatomy, embryology, and cell biology 212, v-115.

Soulika, M., Kaushik, A.-L., Mathieu, B., Lourenço, R., Komisarczuk, A.Z., Romano, S.A., et al. (2016). Diversity in cell motility reveals the dynamic nature of the formation of zebrafish taste sensory organs. Development 143(11), 2012–2024. doi: 10.1242/dev.134817.

Stewart, S., and Stankunas, K. (2012). Limited dedifferentiation provides replacement tissue during zebrafish fin regeneration. Dev Biol 365(2), 339–349. doi: S0012-1606(12)00114-5 [pii] 10.1016/j.ydbio.2012.02.031.

Stylianopoulou, E., Skavdis, G., and Grigoriou, M. (2014). Zinc-Based Fixation for High-Sensitivity In Situ Hybridization: A Nonradioactive Colorimetric Method for the Detection of Rare Transcripts on Tissue Sections. New York: Springer Science+Business Media.

Tachibana, T., Ishizeki, K., Sakakura, Y., and Nawa, T. (1984). Ultrastructural evidence for a possible secretory function of Merkel cells in the barbels of a teleost fish, Cyprinus carpio. Cell and Tissue Research 235(3), 695–697. doi: 10.1007/bf00226971.

Thorimbert, V., Konig, D., Marro, J., Ruggiero, F., and Jaźwińska, A. (2015). Bone morphogenetic protein signaling promotes morphogenesis of blood vessels, wound epidermis, and actinotrichia during fin regeneration in zebrafish. FASEB J 29(10), 4299–4312. doi: 10.1096/fj.15-272955.

Tornini, V.A., and Poss, K.D. (2014). Keeping at arm’s length during regeneration. Dev Cell 29(2), 139–145. doi: 10.1016/j.devcel.2014.04.007.

Tornini, V.A., Thompson, J.D., Allen, R.L., and Poss, K.D. (2017). Live fate-mapping of joint-associated fibroblasts visualizes expansion of cell contributions during zebrafish fin regeneration. Development 144(16), 2889–2895. doi: 10.1242/dev.155655.

Van Trump, W.J., and McHenry, M.J. (2008). The morphology and mechanical sensitivity of lateral line receptors in zebrafish larvae (*Danio rerio*). Journal of Experimental Biology 211(13), 2105–2115. doi: 10.1242/jeb.016204.

von der Emde, G., Mogdans, J., and Kapoor, B.G. (2004). The senses of fish: Adaptations for the reception of natural stimuli. Springer Science+Business Media Dordrecht: Springer Netherlands.

Wehner, D., and Weidinger, G. (2015). Signaling networks organizing regenerative growth of the zebrafish fin. Trends Genet 31(6), 336–343. doi: 10.1016/j.tig.2015.03.012.

Whitear, M. (1992). “Solitary chemosensory cells,” in Fish Chemorecepion, ed. T.J. Hara. 2nd ed (London, UK: Elsevier), 103–125.

Whitfield, T.T., Granato, M., van Eeden, F.J., Schach, U., Brand, M., Furutani-Seiki, M., et al. (1996). Mutations affecting development of the zebrafish inner ear and lateral line. Development 123(1), 241–254.

Williams Iv, R., Neubarth, N., and Hale, M.E. (2013). The function of fin rays as proprioceptive sensors in fish. Nature Communications 4, 1729. doi: 10.1038/ncomms2751.

Williams, R., and Hale, M.E. (2015). Fin ray sensation participates in the generation of normal fin movement in the hovering behavior of the bluegill sunfish (*Lepomis macrochirus*). The Journal of Experimental Biology 218(21), 3435–3447. doi: 10.1242/jeb.123638.

Woo, S.-H., Lumpkin, E.A., and Patapoutian, A. (2015). Merkel cells and neurons keep in touch. Trends in Cell Biology 25(2), 74–81. doi: 10.1016/j.tcb.2014.10.003.

Zaccone, G. (1986). Neuron-specific enolase and serotonin in the Merkel cells of conger-eel (Conger conger) epidermis. Histochemistry 85(1), 29–34. doi: 10.1007/bf00508650.

Zaccone, G., Fasulo, S., and Ainis, L. (1994). Distribution patterns of the paraneuronal endocrine cells in the skin, gills and the airways of fishes as determined by immunohistochemical and histological methods. The Histochemical Journal 26(8), 609–629. doi: 10.1007/bf00158286.

Zachar, P.C., and Jonz, M.G. (2012). Confocal imaging of Merkel-like basal cells in the taste buds of zebrafish. Acta Histochemica 114(2), 101–115. doi: 10.1016/j.acthis.2011.03.006.

Żmijewski, M.A., Sweatman, T.W., and Slominski, A.T. (2009). The melatonin-producing system is fully functional in retinal pigment epithelium (ARPE-19). Molecular and Cellular Endocrinology 307(1-2), 211–216. doi: 10.1016/j.mce.2009.04.010.

