## Supplementary figures and images for "Distribution and restoration of serotonin-immunoreactive paraneuronal cells during caudal fin regeneration in zebrafish"

### Supplementary figure

Supplementary figure

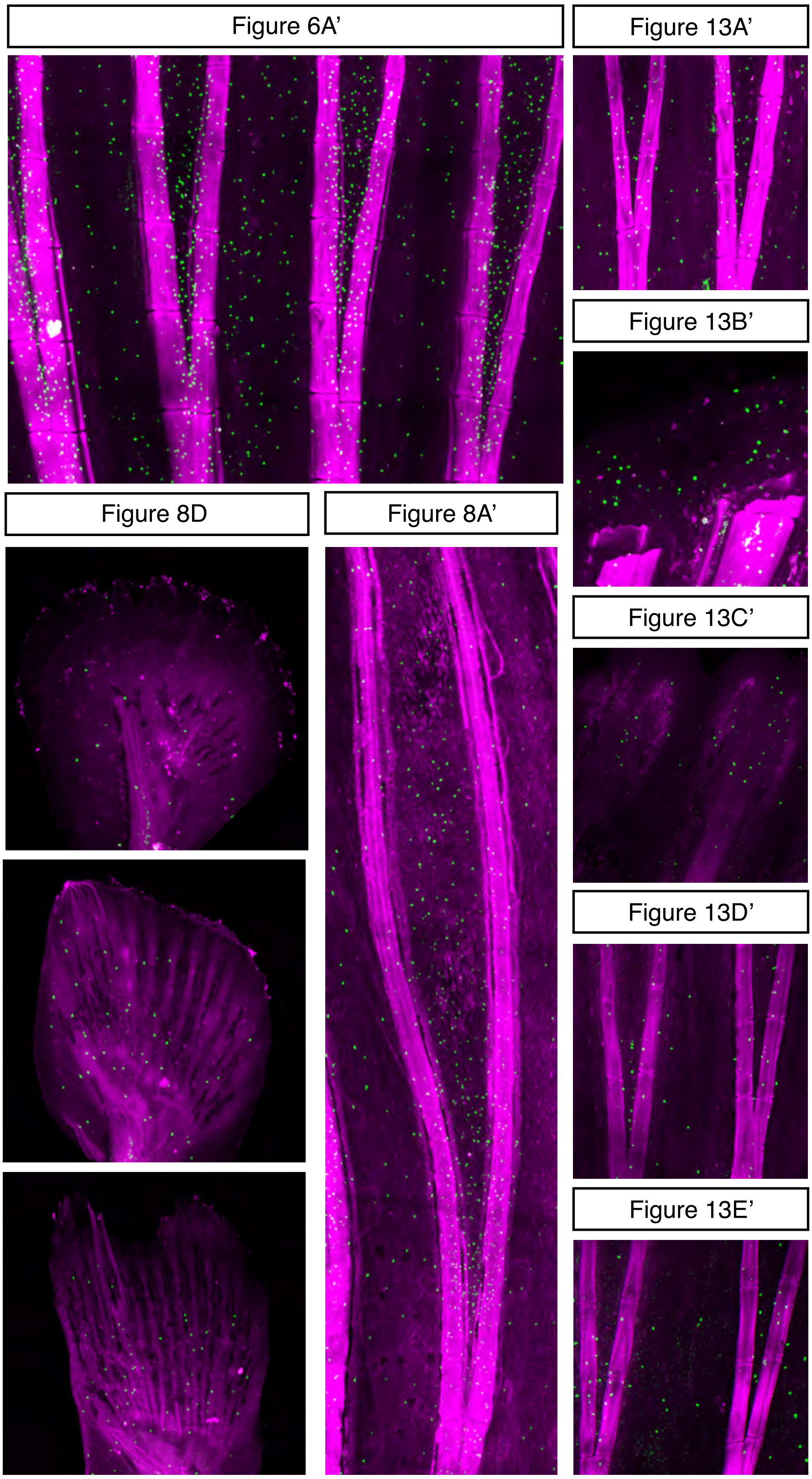
